# Early hematopoietic fate decisions are independent of bone marrow niche

**DOI:** 10.64898/2026.10.20.760390

**Authors:** Christian A. Reardon-Lochbaum, Paolo Cadinu, Hao Zhang, Alan B. Cantor, Jeffrey R. Moffitt

## Abstract

Hematopoietic stem cells (HSCs) reside in specialized bone marrow microenvironments, termed niches, which regulate self-renewal and differentiation through local support programs. Although multiple anatomically distinct niches with unique cellular compositions have been described, it remains unclear to what degree these niches differentially shape HSC fate. In parallel, hematopoietic output is modulated with age, yet it remains unclear if this shift in HSC fate is driven by niche-specific modifications. To address these questions, we applied Multiplexed Error Robust Fluorescence in situ Hybridization to jointly define the cellular and spatial organization of the femoral marrow in young and aged mice. We find that while HSC niches are anatomically and compositionally distinct, stem cell lineage priming is not strongly associated with niche identity consistent with our demonstration that fate-shaping signals are provided redundantly across niches. Moreover, we find that while aging remodels the cellular composition of the marrow, reshapes cellular expression, and alters hematopoietic differentiation, HSC niches are preserved, and HSC fate remains niche agnostic. Together these findings separate anatomical niche identity from early HSC fate decisions and are consistent with a model in which hematopoietic support is distributed across different niche locations rather than partitioned among discrete sites.

## Introduction

Bone marrow (BM) maintains blood homeostasis by balancing hematopoietic stem cell (HSC) self-renewal with the continuous generation of lymphoid, myeloid, megakaryocyte, and erythroid lineage cells^1–3^. HSCs initiate differentiation through a continuum of multipotent and lineage-primed progenitor states, including multipotent progenitors (MPPs) and progenitors biased toward lymphoid, myeloid, or megakaryocyte– erythroid fates. Collectively termed hematopoietic stem and progenitor cells (HSPCs), these cells retain multilineage or restricted multilineage potential before irreversible commitment to a single hematopoietic lineage^4,5^. This process is shaped by both cell-intrinsic differentiation programs—coordinated by transcription factors—and cell-extrinsic support programs—characterized by interactions between receptors on HSPCs and ligands produced by surrounding cells^6,7^. Mesenchymal stromal cells (MSCs)^8,9^, endothelial cells^10,11^, osteolineage cells^12,13^, and megakaryocytes (MKs)^14,15^, among others, have each been observed to provide signals that, in part, govern HSC differentiation into lineage-primed progenitors through adhesive cues, chemokines, cytokines, and growth factors.

Both the initial lineage-priming of HSCs and full lineage commitment of lineage-primed progenitors occur in the context of spatially distinct BM anatomical regions, such as the endosteum, the arteriole, and the sinusoid^11,16^. As the cells that provide HSC shaping signals are in different abundance in these anatomical regions, a natural proposal has been that early hematopoietic fate decisions are, at least in part, differentially shaped by a series of unique microenvironments—so called niches—created within these regions, and, as a result, HSPC development is spatially orchestrated within the BM^17–20^. However, the spatial organization of HSC maintenance and development in mouse and human has proven difficult to define, and conflicting proposals have been made: some have argued that HSCs are primarily localized to the arteriolar^21^, endosteal^22^, megakaryocytic^23^, or sinusoidal niches^24^ while more recent work has proposed a preference for multiple niches^25–27^ or no niche preference at all^28,29^. Finally, while the localization of multipotent and lineage-primed progenitor states has received less attention—perhaps due to the evolving definition of the functional divide between HSPC subsets^30–32^—conflicting models have also emerged regarding whether anatomically distinct niches preferentially support specific HSPC states^16,33,34^. Thus, despite decades of work, the broad HSPC niche model remains operationally ill-defined, with competing anatomical models that have yet to be reconciled into a single spatial framework for HSPC maintenance and development.

In parallel, BM hematopoietic output is shaped by age, physiological demand, and disease state^35–37^, with shifts in hematopoietic output occurring alongside global remodeling of BM structure and transcriptional remodeling of HSPCs. Aging provides a striking example: total HSPC populations expand, yet this accumulation of progenitors does not translate proportionally into productive hematopoiesis^38,39^. Instead, output becomes increasingly biased toward myeloid and megakaryocytic lineages at the expense of lymphoid and erythroid production^40^. Alongside these shifts, the BM undergoes broad molecular and structural remodeling, with an upregulation of inflammation associated pathways in a process termed inflammaging^41–43^. It has been proposed that this shift in hematopoietic output may be due to both cell-intrinsic HSPC alterations as well as reshaped interactions within specific BM niches^29,41–46^. However, age-dependent changes in HSPC localization and niche-specific molecular remodeling remain poorly defined, leaving unresolved the degree to which individual niches regulate age-related changes in hematopoietic output.

Here, we define the spatial orchestration of murine HSPC maintenance and fate decisions using Multiplexed Error Robust Fluorescence in situ Hybridization (MERFISH)^47–50^. We profile the expression of 1,021 RNAs in 2.1 million cells from 34 femoral cross sections across 8-, 40- and 90-week-old mice. We define HSPC subsets as well as the full complement of mesenchymal, endothelial, and osteolineage populations and register these cells in the context of endosteal, sinusoidal, megakaryocytic, and central marrow niches. We find that, within each niche, HSCs, MPPs, and all lineage-primed progenitors are distributed nearly uniformly, indicating that hematopoietic lineage priming is not strongly correlated to niche location. We reconcile this observation with the niche specific cell-type expression of individual HSPC fate-shaping factors by showing that different populations of cells provide the same signals in different niches, revealing that HSPC support is redundant across niches. We find extensive molecular and cellular remodeling in the aging BM, with a global increase in HSPC abundance, a redistribution in the frequency of lineage-primed progenitors, and accumulating transcriptional changes. Yet, despite these broad aging-related changes, we again find that HSPCs show no niche preference and that niche-HSPC interactions are remodeled in a way that preserves the broad signal redundancy across niches. Our results contextualize recent single-cell RNA sequencing (scRNAseq) data and multiplexed immunofluorescence in both mouse and human, and collectively our work supports a model in which the control of HSPC maintenance and fate is not a property of individual BM niches but of the BM as a whole.

## Results

### Mapping the murine femoral bone marrow with MERFISH

To establish a spatial framework for interrogating the role of BM niches in HSPC maintenance and fate, we set out to leverage MERFISH^47–50^ to chart the cellular and spatial structure of the murine BM. However, given the unusually high lability of BM RNA^51–54^, we first evaluated different tissue-preservation strategies for their ability to maintain intact RNA during tissue processing. We observed poor RNA detection and cell-type definition using a wide range of standard fresh- and fixed-frozen protocols^55^ (Supplementary Fig. 1a-o), likely consistent with the remarkable rapidity of RNA degradation post-mortem (Supplementary Fig. 1p,q). To address this challenge, we adopted our recently developed RNA preservation strategy—Rapid Inhibition with Permanent Inactivation of Nucleases (RIPIN)—in which the tissue is rapidly submerged in concentrated cesium sulfate—a paraformaldehyde-(PFA)-fixation compatible analog of the potent RNase inhibitor, RNAlater—to rapidly stabilize RNAs (Methods)^56^. Indeed, we found that murine BM RNA was stable in this solution (Supplementary Fig. 1r), and, importantly, that tissues fixed within this solution produced high-quality MERFISH measurements (Supplementary Fig. 1s-u).

We next designed a 1,021-gene MERFISH panel to capture cell identity, differentiation state, and hematopoietic support programs. The panel included markers of hematopoietic and mesenchymal lineages, large portions of the Hippo-Yap, RAS, HIF, Notch, Wnt, TGFβ, TNF, and other signaling pathways, and broad coverage of integrins, cytokines, chemokines, and their receptors (Methods). With this library, we imaged 34 femoral cross-sections from 8-, 40-, and 90-week-old male mice as well as 8-week-old female mice, comprising 137 million individual RNA molecules (Fig. 1a). Marker transcripts revealed the expected gross anatomy of femoral BM and surrounding bone, including dense hematopoietic regions, bone-associated surfaces, and stromal structures (Fig. 1b). Moreover, global expression profiles were reproducible between biological replicates and correlated with bulk RNA sequencing of extracted mouse BM, supporting the quality of the measurements (Supplementary Fig. 2a-c). RNA molecules were then assigned to single cells using a multi-step segmentation strategy, in which cell boundaries were identified via Cellpose^57^ trained on a nuclear and a cell boundary stain, RNAs were assigned to these cells, and then the RNA-cell assignment was refined using RNA identity and distribution with Baysor^58^ (Fig. 1c; Methods). Spurious segmentation boundaries and small cell fragments were removed, leaving 2,165,026 cells.

**Figure 1.**
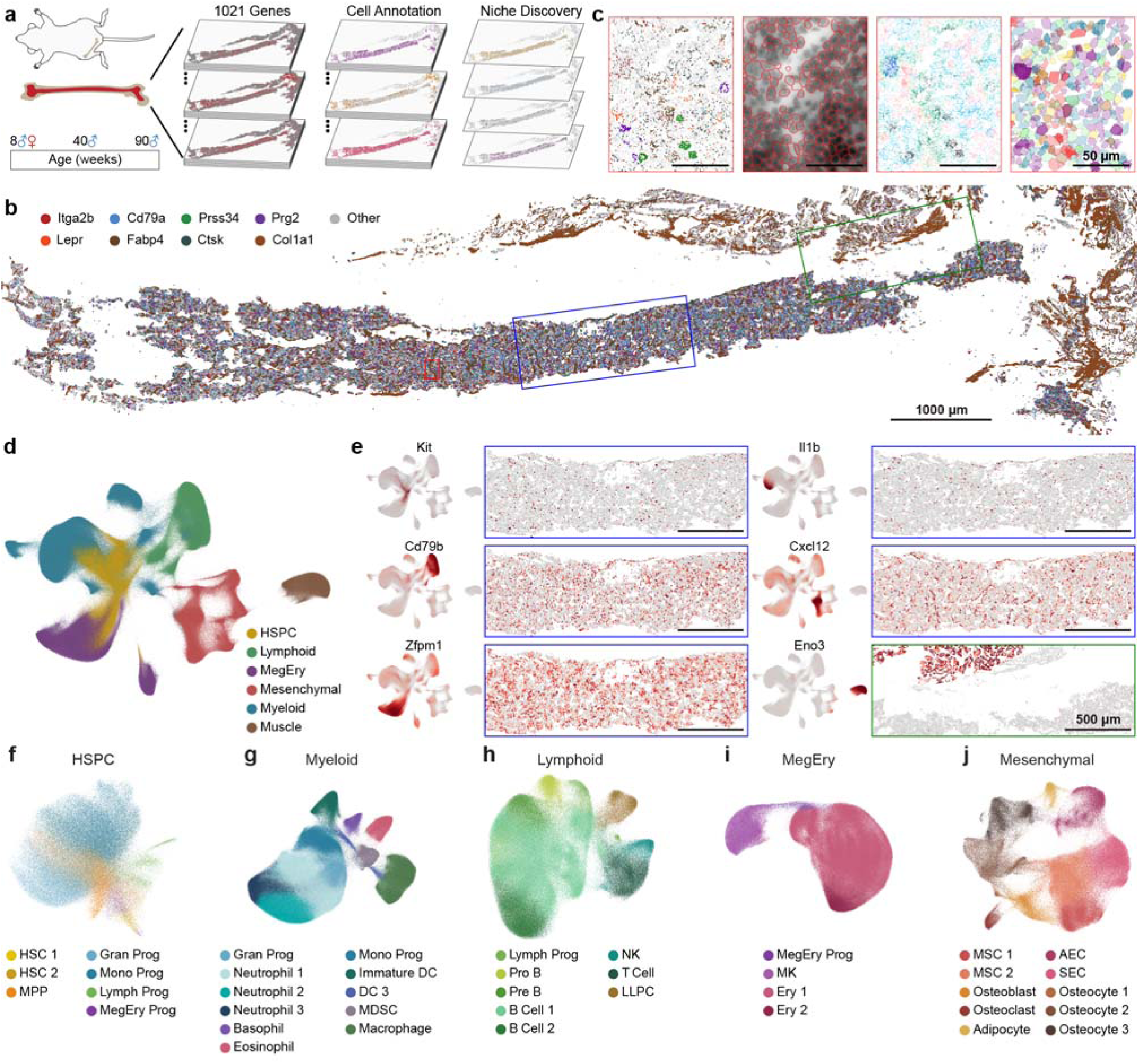
MERFISH resolves the cellular and spatial organization of mouse femoral bone marrow. **a**, Experimental design for spatial profiling of murine femoral bone marrow across age and sex with MERFISH. **b**, Representative femoral section from an 8-week-old male mouse showing the spatial distribution of 1,021 RNAs (gray) with 8 specific RNAs highlighted (color). Scale bar: 1000 µm. **c**, Representative zoom-in on (b) illustrating the cell segmentation pipeline, highlighting RNA identity (left), intermediate cell boundaries superimposed on a DAPI image (left middle), RNAs colored by final cell assignment (right middle), and final cell boundaries colored by cell type (right). Scale bar: 50 µm. **d**, UMAP of all cells colored by major cell class. **e**, UMAP (left) and matched spatial expression (right) for representative HSPC, lymphoid, megakaryocytic-erythroid (MegEry), myeloid, mesenchymal, and muscle markers. The spatial expression shows cell centroids colored by expression of the listed gene for the similarly colored regions in (b). The UMAP is colored by the expression of the listed gene. Expression is normalized to the maximum observed value. Scale bar: 500 µm. **f**-**j**, Compartment-specific UMAPs resolving HSPC (f), myeloid (g), lymphoid (h), megakaryocytic-erythroid (i), and mesenchymal/stromal (j) cell states.

To identify cell types and states, we performed two tiers of clustering which partitioned major cell classes—hematopoietic stem and progenitor cells (HSPCs), myeloid cells, lymphoid cells, megakaryocyte and erythroid cells (MegEry), mesenchymal or stromal cells, and skeletal muscle cells—into 37 transcriptionally distinct populations (Fig. 1d-j). Marker expression and spatial location supported the labels we assigned to these populations (Fig. 1e), which were further supported by high marker purity (Supplementary Fig. 2d) and strong co-integration with published scRNAseq of the murine BM^59^ (Supplementary Fig. 2e,f). Cells from both sexes and all ages co-integrated, indicating that none of these populations was shaped solely by sex or age (Supplementary Fig. 2g,h).

### MERFISH resolves hematopoietic and mesenchymal differentiation states

A central requirement for profiling HSPC niches is resolving both the differentiation state of the HSPC and the identity of the surrounding cells that make up its local environment. Thus, we first examined our resolution in the HSPC compartment (Fig. 1f; Supplementary Fig. 3). We identified two HSC subsets, a multipotent progenitor (MPP), and four lineage-primed progenitors, associated with monocytes (Mono Prog), granulocytes (Gran Prog), lymphoid cells (Lymph Prog), and the megakaryocytic-erythrocytic lineage (MegEry Prog; Fig. 1f), with the identity and developmental relationship between these populations supported by pseudotime, lineage scores, and marker gene expression (Supplementary Fig. 3a-d). Moreover, differential expression between pairs of these populations revealed the expected shifts along the continuous HSPC lineage development (Supplementary Fig. 3e,f)^5,60–62^. For example, the two HSC subsets, HSC 1 and HSC 2, both expressed stem-associated programs but HSC 2 expressed higher levels of Cd34, Flt3, Myc, and Tgfbr2, relative to HSC 1, consistent with a transition away from quiescence^30,31^. MPPs showed further loss of stem-associated genes, including Kit and Slamf1, and were followed by the progressive acquisition of lineage-specific programs. Consistent with known differentiation trajectories, MegEry Prog showed a rebound in the HSC-like-program relative to MPPs^63,64^ and increased expression of markers of this lineage, e.g., Runx1 and Vwf; Lymph Prog were upregulated in Dntt and Il7r; Mono Prog showed increased levels of Irf8, Il6ra, and Cebpa; and Gran Prog had increased levels of Cebpa, Cebpe, and Gfi1.

We next turned our attention to the remaining hematopoietic and stromal compartments. Among these cell classes, we resolved the expected cellular diversity (Fig. 1g–i; Supplementary Fig. 4). Within myeloid cells, we identified a progression of increasingly mature neutrophil states, a diversity of immature and BM resident mature dendritic cells (immature DC and DC 3)^65^, macrophages, and myeloid derived suppressor cells (MDSC). Similarly, among lymphoid cells, we identified natural killer (NK) cells, T cells, and the continued maturation of B cells from Pro B, to Pre B, and then to B Cells 1 and 2. We also identified a population of long-lived plasma cells (LLPCs), which are recruited back to the BM from circulation^66^. Finally, we also observed the branching of the MegEry Prog to form mature MKs and two states of erythrocyte development (Ery 1 and 2).

Mesenchymal cells are particularly important components of HSPC niches, as they can provide HSC support and fate-shaping factors^8,9^. Importantly, we identified a diversity of mesenchymal and stromal cells, with populations spanning the known transcriptional continuum from mesenchymal progenitor states to adipocytic and osteolineage differentiation^59^. These populations included multipotent mesenchymal stromal cells (MSC 1 and 2), osteoblasts, osteoclasts, adipocytes, and three osteocyte states (Fig. 1j; Supplementary Fig. 4d). We also identified both sinusoidal and arteriolar endothelial cells (SEC and AEC, respectively; Fig. 1j; Supplementary Fig. 4d). Notably, MSC 1 and 2 were divided by genes associated with an established MSC activation axis^67^. While both expressed canonical hematopoietic-supportive stromal programs, e.g., Lepr and Cxcl12, MSC 2 superimposed an immune- and stress-associated program characterized by increased H2-Eb1 and Hmox1 expression (Supplementary Fig. 4d). Although analogous activated MSC states have been described in disease^67^, our work now reveals that they are present in homeostasis as well.

Importantly, our measurements clearly defined a series of populations that have proven challenging to capture with dissociation-based techniques, including eosinophils, basophils, and mature MKs, allowing us to define the potential roles that these cells may play in shaping HSPC fate in different niches. Inclusion of mature MKs is particularly important as these cells have been shown to be important sources of HSPC-fate-shaping factors^14,15^. We, thus, conclude that our atlas provides the cellular resolution necessary to both define the subtle divisions within early HSPC lineage development as well as the wide diversity of cells that comprise the different BM niches proposed to shape this development.

### Bone marrow is organized into broad anatomical regions with recurrent spatial motifs

We next sought to define the spatial organization of the BM, focusing first on young adult (8-week-old) male mice. To separate different gross anatomical structures in the BM, we leveraged a neighborhood analysis to classify tissue features based on the local composition of cells^68^ (Methods). This analysis cleanly partitioned our sections into four major neighborhoods: marrow, bone, arteriole, and muscle (Fig. 2a-d). To then define the cellular-scale structure of the BM, we selected cells that were assigned to either the bone, marrow, or arteriole neighborhoods and calculated a local spatial enrichment between cell pairs (Fig. 2e; Methods).

**Figure 2.**
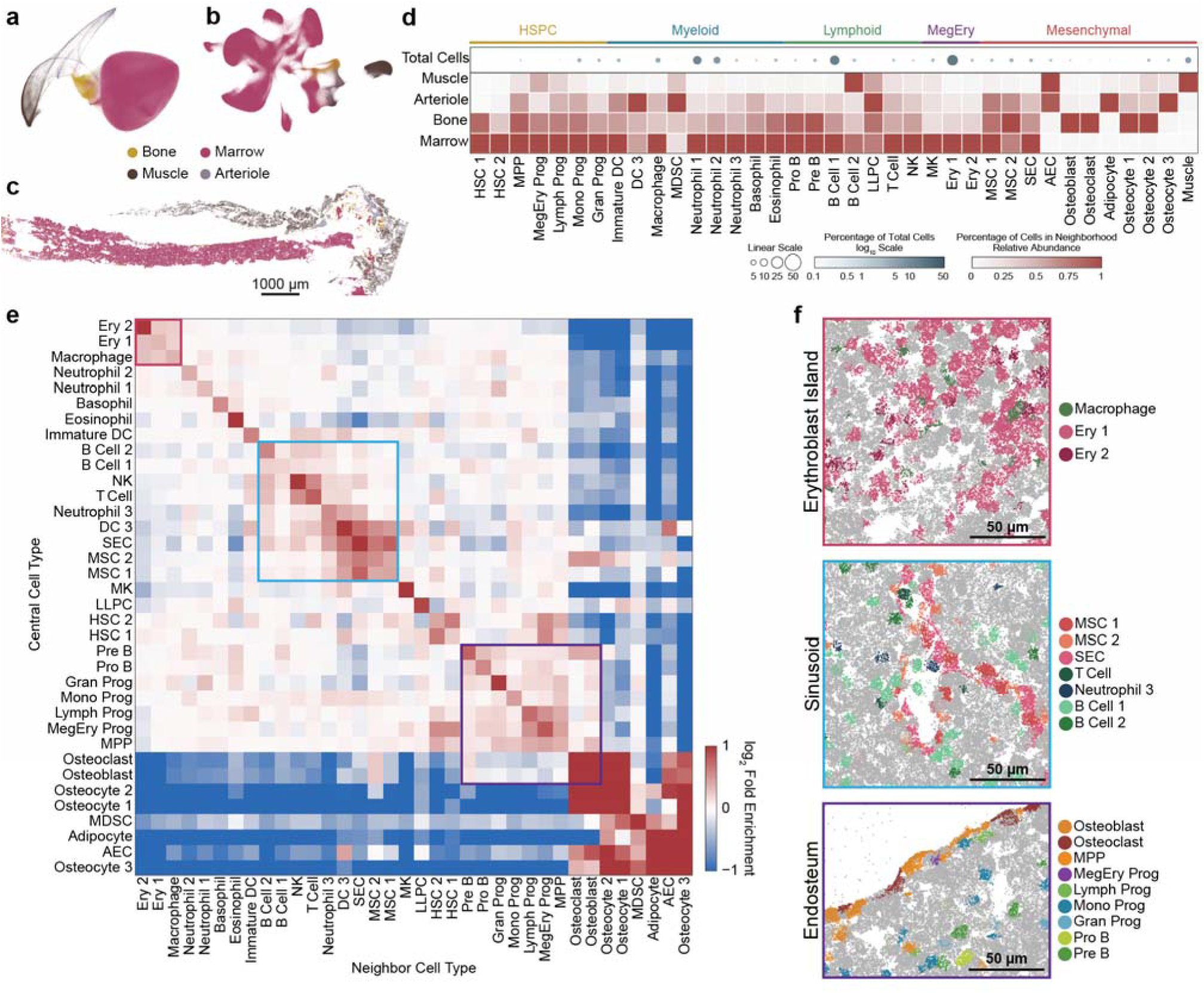
Mouse bone marrow is locally organized into spatial motifs. **a**,**b**, UMAP representation of cellular neighborhoods (a) or cell type (b) colored by the four defined tissue neighborhoods. **c**, Representative femoral section from an 8-week-old male mouse with cell centroids colored by neighborhood assignment. Scale bar: 1000 µm. **d**, Relative abundance of each annotated cell type across the four neighborhood classes. Circles indicate the overall abundance of each cell type in the dataset, with size and color both indicating abundance on different scales. **e**, Pairwise enrichment of all neighboring cells within a 25 µm radius of each central cell. Colored boxes highlight Erythroblast Island (red), Sinusoid (blue), and Endosteum (purple) spatial motifs. **f**, Spatial distribution of RNAs colored by the cell type of origin or gray if not associated with a listed cell type. Representative cell types for each spatial feature are shown. Scale bars: 50 µm.

This analysis revealed a diversity of local spatial enrichments. First, there was a clear partitioning between the cellular structure of the bone, in which we observed pairwise enrichments between cells associated with bone maintenance—osteocytes, osteoblasts, and osteoclasts. As expected, bone structures were the dominant sites of AECs, as arterioles are localized to the endosteum^20^. MDSCs were also strongly associated with osteocyte microenvironments, confirming their role in regulation of bone inflammation^69^ and their known restriction to bone environments in young mice.

Although pairwise enrichments within the marrow were relatively modest, several major spatial modules emerged. The most prominent example was the enrichment of macrophages with Ery 1 and 2 populations, recapitulating the erythroblastic island in which developing erythroid cells mature around a central macrophage^70^ (Fig. 2e,f). We also observed a clear sinusoidal environment, with strong pairwise enrichment between SECs and both MSC states (Fig. 2e,f). Interestingly, a variety of mature immune populations showed stronger enrichment with these sinusoidal associated cells, including DC 3, neutrophil 3, T cells, and NK cells. This local enrichment of mature immune cells at the sinusoids may be important for immune surveillance and trafficking^65,71,72^. Finally, we observed an endosteal environment, which contained the expected osteoblasts and osteoclasts as well as a slight enrichment in early progenitor HSPC states and early states in B cell differentiation (Fig. 2e,f), consistent with previously described interactions between osteolineage cells and HSPCs in the endosteal niche^18^ and the specific osteolineage-support for committed pre and pro B cells^73,74^.

Together, these spatial enrichments demonstrate that BM is organized into broad anatomical regions with recurrent local microenvironments. Most hematopoietic spatial enrichments were modest, indicating that individual cell populations participate in multiple local environments rather than occupying exclusive anatomical territories. Importantly, HSPCs were similarly broadly distributed across the marrow, with only modest enrichment near canonical niche-associated populations. Thus, our data suggest that HSPCs are not confined to a single anatomical niche.

### Sexual dimorphisms in young bone marrow are limited

We next examined the sexual dimorphisms present in the BM from 8-week-old male and female mice. Overall, we found that the cellular composition was highly similar between sexes, with only modest differences across HSPC, myeloid, lymphoid, megakaryocytic-erythroid, and mesenchymal populations (Supplementary Fig. 5a-b). We also found the transcriptional differences were similarly modest. Only three cell types—B cell 1, MK, and MSC 1—had any gene with a statistically significant difference in expression greater than 2-fold (Supplementary Fig. 5c). Of these cell types, the strongest sex-associated differences were found within MSC 1, which is consistent with recent reports of sexually dimorphic stromal programs^75,76^. Finally, we found that the overall structure of young female and male BM was essentially indistinguishable. As expected, all four neighborhoods could be found in both sexes, and only a small number of pairwise cell type enrichments showed a statistically significant difference, although all were small (Supplementary Fig. 5d,e). Thus, we conclude that there are limited sexual dimorphisms in the cellular organization of male and female mice. As such, we combined male and female 8-week-old mice for all subsequent analysis.

### HSPC niche occupancy does not align with early hematopoietic differentiation trajectory

Having established that HSPCs are distributed across multiple anatomical environments within mouse BM, we next sought to define the niches occupied by individual HSPCs and determine whether particular HSPC states preferentially associate with anatomically distinct niches. To this end, we explored the local transcriptional and cellular microenvironment around each HSPC (Methods). We found that local transcriptional profiles could be classified into four distinct HSPC niches—central marrow, sinusoidal, megakaryocytic, and endosteal—distinguished based on their transcriptional profile, cellular composition, and anatomical location (Fig. 3a-c; Supplementary Fig. 6a). While these niches broadly corresponded to those previously described^19^, we noted that this analysis did not identify an arteriole niche. The arterioles connect the central artery in the BM to the endosteum and ultimately the sinusoids and are rare, which may explain both why we did not identify a unique niche and why the few AECs we observe within the marrow were associated with these two niches.

**Figure 3.**
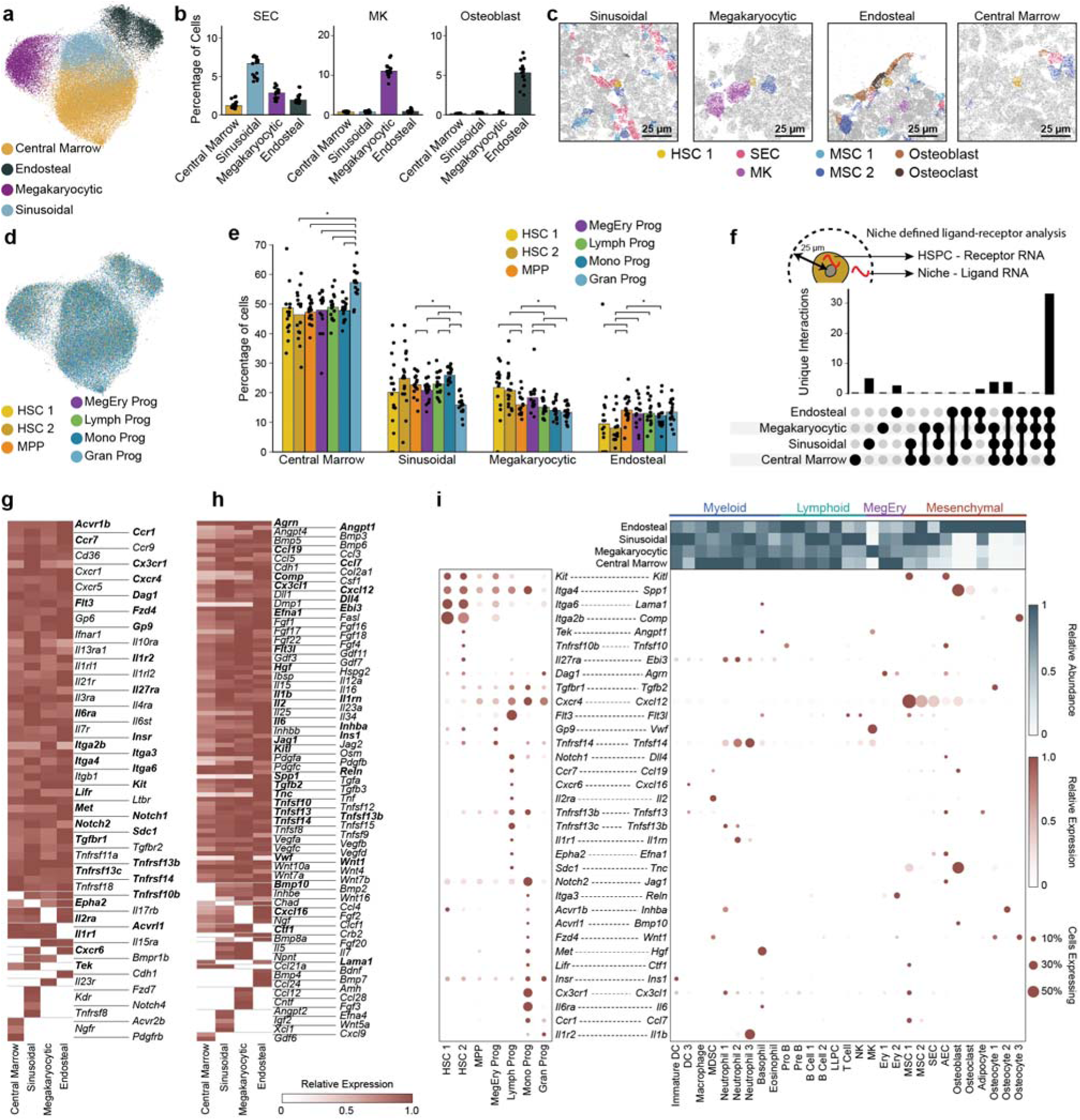
Anatomically distinct HSPC niches contain similar progenitor state proportions and overlapping signaling programs. **a,** UMAP of the gene expression in a 25-µm window of each HSPC colored by niche assignment. **b,** Abundance of niche-defining cell populations across niche classes, illustrated for sinusoidal endothelial cells (SEC), megakaryocytes (MK), and osteoblasts. Bars: mean. Markers: individual bones. **c,** Spatial distribution of RNAs colored by listed cell types or gray for all other RNAs for representative instances of each spatial niche. Scale bars: 25 µm. **d,** UMAP as in (a) but colored by the central HSPC state. **e,** Fraction of each HSPC state assigned to each niche. Bars: mean. Markers: individual bones. Brackets highlight pairwise comparisons for which the FDR-corrected P value is less than 0.05. **f,** Number of spatially restricted receptor-ligand interactions with the receptor provided by an HSPC and the ligand from the surrounding niche. **g,h,** Relative expression of candidate HSPC receptor genes (g) and corresponding niche ligand genes (h) across niche classes. **i,** Integrated view of selected ligand-receptor pairs showing receptor expression across HSPC states (left), the relative abundance of neighboring cell types across niches (top), and ligand expression across neighboring cell populations (right). Circle size denotes the fraction of cells expressing a gene and color denotes relative expression. Circles are absent if cell-specific expression was below the false discovery threshold. Receptor-ligand pairs shown in (i) are bolded in (g) and (h).

We then examined the frequency with which different HSPC states were found within each niche (Fig. 3d,e). Our measurements identified a few statistically significant niche preferences. For example, relative to the lineage-primed states and MPPs, HSC 1 and 2 were found more frequently in the megakaryocytic niche and less frequently in the endosteal niche. Similarly, Gran Prog were found more frequently in the central marrow than other HSPC states. However, in each of these cases, the difference in frequency was modest. And while the total fraction of each HSPC state associated with a niche largely tracked the abundance of that niche within the BM, each of the different HSPC states were found at very similar frequencies within a given niche (Fig. 3d,e). These results support a model in which niche identity does not play a major role in shaping early HSPC fate decisions.

Supporting the lack of a strong niche compartmentalization for any specific HSPC subtype, we examined the enrichment of individual HSPC subsets next to cell types that define the sinusoidal, endosteal, and megakaryocytic niches, as well as the arterioles, even though our analysis above did not define an arteriolar niche. In all cases, no pairwise enrichment or depletion was greater than 2-fold (Supplementary Fig. 6b-c), supporting our observation that there is no definitive partitioning of any HSPC subset to a single or limited subset of niches. We then reanalyzed multiplexed proteomic imaging of mouse femoral BM^77^ and found that even with a more limited definition of HSPC states, these data provided independent support for our observations by revealing no niche preference (Supplementary Fig. 6d-g).

We next asked whether niche location was associated with subtle shifts in HSPC transcriptional state. While different HSPC subsets showed substantial numbers of differentially expressed genes (Supplementary Fig. 3f) and the niches themselves showed clear differential gene expression (Supplementary Fig. 6h), HSPCs showed essentially no differential expression when found in different niches (Supplementary Fig. 6i,j). Thus, we conclude that HSPC subsets neither have a strong preference for distinct niches nor are differentially modulated by individual niches in their gene expression, at least among the genes we measured. To explore whether a similar spatial organization might be present in human BM, we analyzed a human multiplexed proteomic imaging dataset^27^ (Supplementary Fig. 7a,b). We constructed a radial cell-neighborhood matrix analogous to that used to define niches in our mouse MERFISH dataset and identified five HSPC-associated microenvironments: Central Marrow, Megakaryocytic, Perivascular, Adipo-MSC, and Thy1+ MSC niches (Supplementary Fig. 7c-e). These broadly recapitulated the major niche classes identified in mouse, although the human Perivascular niche encompassed sinusoidal, arteriolar, and endosteal regions. The Adipo-MSC and Thy1+ MSC niches did not have direct counterparts in the mouse dataset, consistent with greater stromal heterogeneity in human BM. Despite these anatomical differences, HSPC populations were distributed similarly across niches, with no clear enrichment of specific lineage trajectories within individual niches (Supplementary Fig. 7f). Together, these cross-species analyses suggest that the niche-agnostic fate of HSPCs seen in mouse is also present in humans.

### Spatially distinct niches provide overlapping hematopoietic support programs

An argument in favor of the niche-centric model of HSPC fate is that specific molecular signals are associated with HSC quiescence, maintenance, and developmental bias towards individual hematopoietic lineages, and, critically, these signals are provided by cell types that are not uniformly found across different niches^6–15^. Thus, we next asked how HSPC fate can be niche agnostic in the presence of niche-enriched, cell-type-specific signals.

To this end, we examined the differential signaling between HSPCs and cells that comprise different niches, focusing on receptors expressed by HSPC subsets and ligands expressed by cells in their vicinity (Methods). Supporting our analysis, we recovered a diversity of established interactions known to drive HSPC maintenance^6,8,21,78^. For example, Kit was broadly expressed across HSPCs while its ligand, Kitl, was produced by MSCs, AECs, and osteoblasts (Supplementary Fig. 8a). Similarly, we identified Flt3l-Flt3 interactions, known to serve multiple roles in the progression from HSC to lineage primed progenitor states, between HSC 2, MPPs, and all lineage-primed progenitors and DCs, Neutrophils, Macrophages, B cells, and MSCs (Supplementary Fig. 8b). We also identified known factors of lineage-priming. For example, Jag1, which supports lymphoid development via Notch2^31^, was expressed by AECs and SECs (Supplementary Fig. 8c). Similarly, Il1b, which supports myeloid differentiation via Il1r1^79^, was expressed by Neutrophils, DC 3, NK, and MSC 2 (Supplementary Fig. 8d).

We next interrogated the anatomical restriction of specific signals provided to HSPCs. Strikingly, most candidate ligand–receptor interactions were shared among multiple niches with the majority of cognate ligand-receptor pairs expressed in all four niches (Fig. 3f-h; Supplementary Fig. 8a-d). Resolving this shared landscape across niche ligand expression and HSPC receptor state showed that this overlap arose from broad availability on both sides of the interaction: receptors were generally represented across HSPCs in all niches while their cognate ligands were similarly expressed within most niches (Fig. 3g,h). Importantly, this broad niche-redundant signaling was not indicative of broad cellular expression of these signals. Rather, many receptors and their cognate ligands were expressed only by a small subset of cell types. However, these cells or different combinations of these cells were found in each niche (Fig. 3i). Thus, we conclude that there is a surprising degree of signaling redundancy across anatomically distinct niches, such that niches with different cell type compositions can, nonetheless, provide highly similar HSPC-supportive signaling environments.

This shared signaling architecture encompassed both established regulators of HSPC biology and less-characterized candidate niche signals (Fig. 3i). In addition to well-described interactions including Kitl–Kit, Flt3l–Flt3, Jag1–Notch2, and Il1b–Il1r1 (Supplementary Fig. 8a–d), we identified broadly distributed candidate interactions such as Tnfsf14–Ltbr, Cx3cl1–Cx3cr1, and Agrn–Dag1, implicating immune-regulatory, chemokine, and extracellular-matrix-associated signaling in HSPC microenvironments^80–82^. Despite the predominance of signals shared across niches, a smaller subset—including Efna1–Epha2, Il2– Il2ra, Tnfsf10–Tnfrsf10b, and Cxcl16–Cxcr6—were restricted to particular niches (Fig. 3g–i), indicating that overlapping core support may be accompanied by a more limited layer of niche-specific signaling that may provide potential mechanisms through which individual niches could fine-tune HSPC state without imposing distinct lineage programs. Finally, analysis of independent mouse and human BM scRNAseq datasets reproduced the cell-type-specific expression patterns of these niche factors (Supplementary Fig. 8e,f), providing an additional layer of support for the cell-type signaling relationships identified by MERFISH.

Together, these findings reveal a distinction between the anatomical identity of an HSPC niche and the unique HSPC-responsive molecular signals available within it. Although central marrow, sinusoidal, megakaryocytic, and endosteal niches contain different cellular compositions and transcriptionally distinct neighboring cells, they appear to have few spatially restricted HSPC fate shaping signals, consistent with the lack of a strong niche preference we observe for HSPC subsets.

### Aging remodels the molecular and cellular landscape of the bone marrow

While our data suggests that niche identity plays little role in HSPC fate in healthy mice, it is possible that individual niches may play specific roles in reshaping hematopoietic fate in response to insult or injury. To this end, aging represents a natural perturbation to hematopoiesis, as age-related changes include a well-established shift away from lymphoid and erythroid production toward myeloid and megakaryocytic output and increased inflammatory signaling^29,83,84^.

To explore how age associated changes to the BM microenvironment might be responsible for age-related changes in hematopoietic output, we compared femoral BM from 8-, 40-, and 90-week-old mice, representing young adult, middle-aged, and geriatric states, respectively (Fig. 4a). All HSPC populations increased markedly in abundance at 40 and 90 weeks (Fig. 4b); however, the relative composition of the expanded HSPC compartment was progressively redistributed, with increased representation of MPPs and Gran Progs at the expense of MegEry, Lymph, and Mono Progs while HSC 1 and 2 populations remained relatively stable (Fig. 4c). These findings indicate that aging expands the HSPC compartment while rebalancing lineage-primed progenitors towards a granulocytic fate.

**Figure 4.**
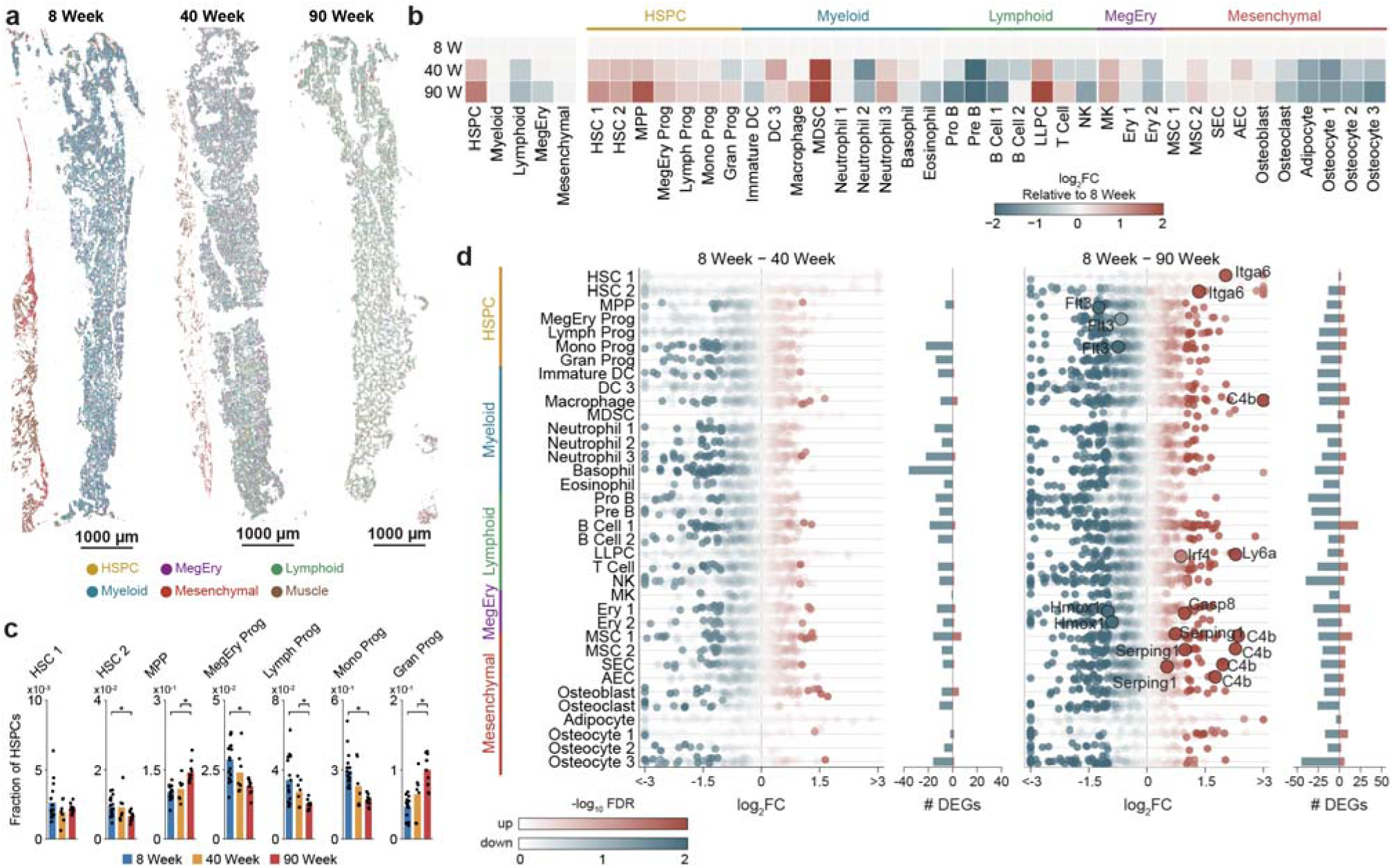
Aging broadly remodels bone marrow and shifts the HSPC compartment toward granulocyte-primed states. **a,** Spatial distribution of cells in representative femoral sections from 8-, 40-, and 90-week-old mice colored by major cell class. Scale bar: 1000 µm. **b,** Abundance of major class and annotated cell states, shown as log_2_ fold change relative to 8-week-old bone marrow. FC: Fold change. **c,** Fraction of total HSPCs per age associated with each HSPC state. Bars: mean. Markers: individual bones. Brackets highlight pairwise comparisons for which the FDR-corrected P value is less than 0.05. **d,** Differential expression for all genes within each cell type for 40-week-old mice (left) or 90-week-old mice (right) relative to 8-week-old mice. Color indicates -log_10_ FDR-corrected P values. The side bars indicate the number of statistically significant up-(red) or down-(blue)-regulated genes as defined by an FDR-corrected P value less than 0.05 and an absolute log_2_ fold change greater than 1. Illustrative genes are highlighted.

We observed similar trends in the transcriptional programs in the HSPC states. At 90 weeks, this broad age-associated program was evident across the HSPC continuum. HSC 1 and HSC 2 showed increased expression of Selp and Itga6, which have been previously associated with impaired hematopoietic function^85–87^ (Fig. 4d; Supplementary Fig. 9a-b). Transcriptional remodeling was exacerbated in more differentiated states. MPPs lost stem-associated genes including Hlf, Meis1, and Angpt1 while inducing Csf2rb, C4b, and Tnfsf15, consistent with an erosion of primitive transcriptional programs and increased capacity to respond to inflammatory cytokines^39,40,46^ (Supplementary Fig. 9c). Strikingly, analogous changes extended into lineage-primed progenitors (Supplementary Fig. 9d-g). Lymph Progs lost Ebf1 and Cd79b—components of the B-lineage specification program^88^—while gaining Ly6a, Il1rl1, and C4b, indicating a relative loss of lymphoid identity accompanied by acquisition of a stress- and inflammatory-associated state^40^. Mono and Gran Progs upregulated Csf2rb, C4b, and Cdk6, suggesting increased cytokine responsiveness with a more proliferative state. Conversely, MegEry Progs increased Itga2b, Selp, and Itga6 together with C4b, consistent with enhanced megakaryocytic-platelet priming superimposed on the similar inflammatory program seen in other HSPCs^40^. Thus, aging does not simply change the relative abundance of HSPC subsets. It transcriptionally reprograms cells across the differentiation continuum, weakening adaptive immune identity while broadly imposing inflammatory features.

At 40 weeks, transcriptional changes were comparatively muted, particularly within HSC 1 and HSC 2, but early remodeling was already evident in MPPs and downstream Mono Prog. These populations showed altered expression of Hk3 together with TGFβ-associated regulators including Tgfbr2 and Smad3, as well as changes in cytokine- and stress-response genes, consistent with early metabolic and signaling remodeling preceding the broader inflammatory program observed at 90 weeks^59,84^. Thus, although substantially attenuated relative to geriatric marrow, a recognizable aging-associated HSPC phenotype was already established by middle age.

The surrounding BM environment also changed with age. The overall abundance of a diversity of mature immune populations and stromal populations were shifted (Fig. 4b). Moreover, many cell populations showed inflammation-associated transcriptional remodeling (Fig. 4d). For example, mature myeloid populations, including neutrophils, DC 3, and macrophages, increasingly expressed cytokine-, complement-, and inflammatory-response programs, as indicated by C4b, Csf2rb, and Serping1 expression, while expanded LLPCs and residual erythroid populations acquired activation- and stress-associated states, marked by Ly6a and Casp8, respectively. Mesenchymal remodeling was particularly pronounced: MSC 1 modestly declined as MSC 2 expanded while stromal populations progressively lost features characteristic of young supportive states and acquired inflammatory programs. We also observed some local remodeling in spatial enrichments, primarily with mesenchymal populations (Supplementary Fig. 9h-j). Nonetheless, the frequency with which HSPCs were found next to other populations was largely conserved between the 90-week- and 8-week-old mice (Supplementary Fig. 9j), indicating that despite a variety of age-related changes to the marrow, HSPCs have a similar spatial distribution.

### Aging remodels signaling similarly across all niches

The relative spatial stability of HSPCs, despite profound remodeling of the BM, prompted us to ask more directly whether aging altered the anatomical niches occupied by HSPCs. Using the same HSPC-centered niche classification applied to young marrow, we found the structures of central marrow, sinusoidal, megakaryocytic, and endosteal niches were largely preserved at both 40 and 90 weeks (Fig. 5a). HSPC differentiation states likewise remained broadly represented across each niche, with no strong age-dependent emergence of lineage-specific spatial segregation (Fig. 5b,c; Supplementary Fig. 10a,b). Thus, age-associated changes in HSPC lineage and hematopoietic output occurred without a corresponding redistribution of HSPC states across niches.

**Figure 5.**
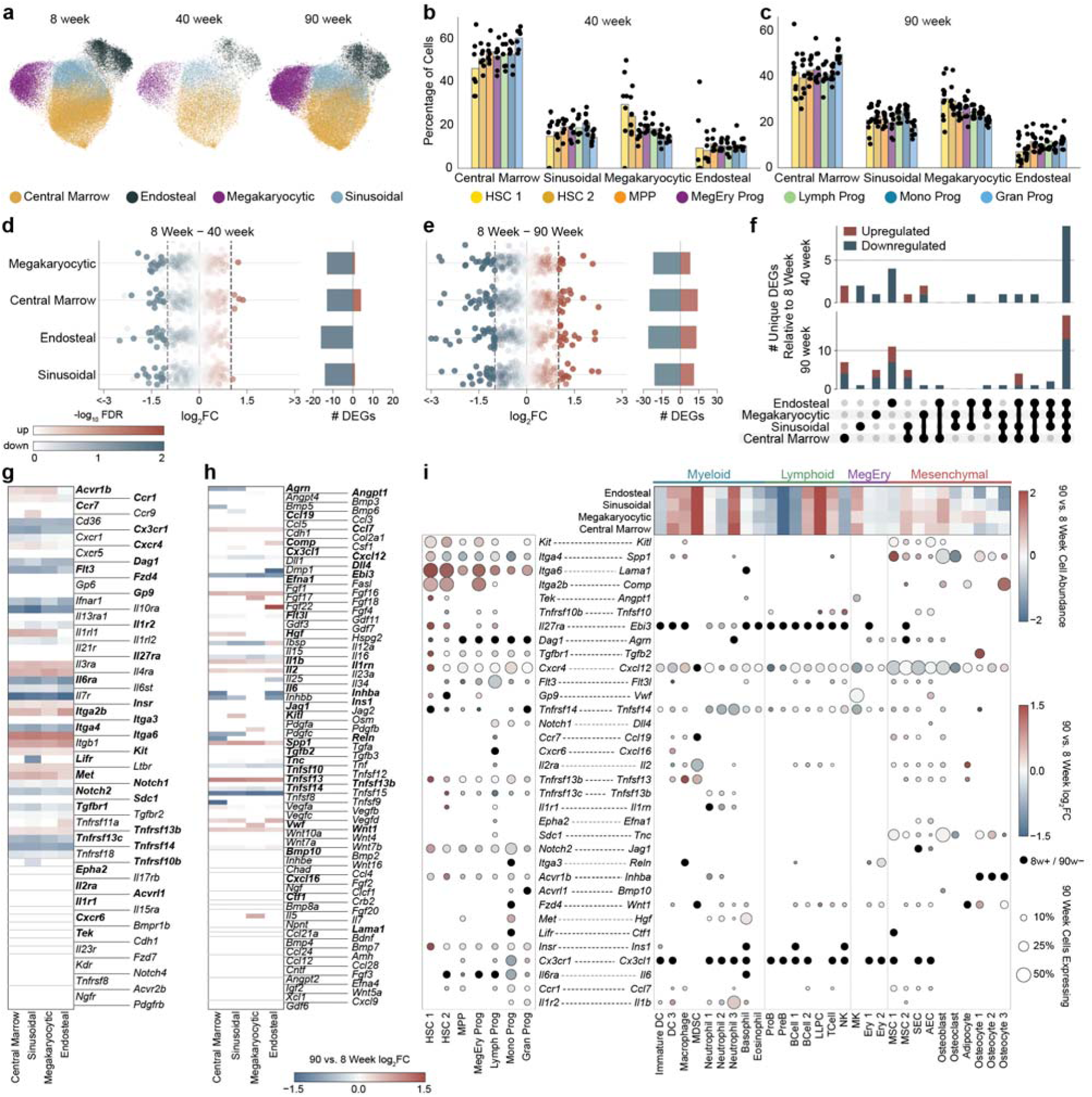
Aging remodels HSPC signaling across niches without generating a spatially restricted aged niche. **a,** UMAP of the gene expression in a 25-µm window of each HSPC colored by niche assignment showing only cells from 8-week-old, 40-week-old, or 90-week-old mice. Cells are colored by niche assignment. **b,c,** Fraction of each HSPC state assigned to each niche for 40-week-old (b) and 90-week-old (c) mice. Bars: mean. Markers: individual bones. **d,e,** Differential expression for all genes within each niche for 40-week-old mice (d) or 90-week-old mice (e) relative to 8-week-old mice. Color indicates -log_10_ FDR-corrected P value. The side bars indicate the number of statistically significant up-(red) or down-(blue)-regulated genes as defined by an FDR-corrected P value less than 0.05 and absolute log_2_ fold change greater than 1. FC: Fold change. **f,** UpSet plots showing the overlap of age-associated upregulated and downregulated genes among niche classes at 40 and 90 weeks relative to 8 weeks. **g,h,** Age-associated log_2_ fold changes in candidate HSPC receptor genes (g) and cognate niche ligand genes (h) across the four niches at 90 versus 8 weeks. **i,** Integrated 90 versus 8 weeks ligand-receptor map, displaying receptor changes across HSPC states (left) and ligand changes across niche cell populations (right), and 90-versus-8-week changes in cell abundance within each niche (top). Dot size indicates the fraction of 90-week cells expressing each gene, color indicates 90-versus-8-week differential expression, and black dots denote genes detected at 8 weeks but not at 90 weeks indicating downregulation. Dots are absent if expression at both 8- and 90-weeks was below the false discovery threshold. Genes plotted in g-i are matched to genes plotted in Fig. 3g-i.

We next asked whether aging reshaped the molecular environment within these niches. Individual niches were clearly remodeled transcriptionally with age (Fig. 5d,e). However, the differentially expressed genes that defined each niche within an age (Supplementary Fig. 10c-e) as well as their magnitude of expression (Supplementary Fig. 10f,g) were conserved. Importantly, genes that were differentially expressed within one niche were most often found differentially expressed across all niches (Fig. 5f), indicating a largely coherent age-related remodeling between niches.

We next asked how this shared remodeling extended to the signaling interactions between niches and HSPCs. Of the receptors expressed on HSPCs, the majority were differentially regulated at 90-weeks but, again, coherently between niches (Fig. 5g). Interestingly, of the ligands expressed within any of these niches, a relatively modest number were differentially expressed (Fig. 5h). The age-related differential expression of both receptors and ligands was cell-type specific (Fig. 5i); however, again, the combination of cell types within different niches largely produced coherent remodeling across niches. Thus, aging altered both the signals available to HSPCs and the receptors through which HSPCs could respond, but these changes were distributed across the marrow rather than confined to one anatomical environment. Collectively, these results suggest that hematopoietic output and HSPC fate can be modulated via cell-type and signal-specific changes but that this modulation is a property of the global BM rather than any specific anatomical niche.

## Discussion

The idea that anatomically distinct niches provide functionally distinct support to orchestrate the maintenance and lineage commitment of HSPCs has long been hypothesized as an organizing principle of the BM^17–20^ yet has remained difficult to fully test. Until the introduction of spatial transcriptomics, no single approach could simultaneously resolve HSPC state, anatomical niche identity, and the expression of the signaling factors mediating interactions occurring within those niches. Unfortunately, application of spatial transcriptomics to murine BM has been limited by the exceptional lability of RNA in this tissue. Here, we overcame the technical challenge of RNA degradation in murine BM, collected a large atlas of the healthy and aging mouse BM with MERFISH, and, with this resource, directly interrogated the long-standing niche-centric hematopoiesis model by identifying individual HSPCs, assigning them to local anatomical niches, and defining the signaling environments associated with each niche.

These measurements reveal that distinct HSPC states are not confined to distinct niches and that anatomically distinct niches provide substantially overlapping HSPC-supportive signaling programs. Thus, our data argue against a model in which early hematopoietic decisions are strictly spatially partitioned among specialized niches. Instead, they support a distributed model in which HSPC support is assembled from multiple cellular sources across different anatomical environments, allowing similar hematopoietic states and outputs to be maintained across multiple niches. Importantly, aging provides an additional test of this framework: even as HSPC composition and hematopoietic output shift dramatically, distinct anatomical niches remodel in parallel rather than providing diverging support for lineage-specific states.

Indeed, recent imaging work is consistent with this distributed support model^26,29^. For example, imaging of genetically labeled functional long-term HSCs and key niche-associated cell populations revealed that adult murine long-term HSCs are found in many different anatomically distinct niches at a frequency indistinguishable from random chance^28^. Similarly, as we show here, recent multiplexed immunofluorescence imaging of both mouse and human BM are also consistent with limited niche preference^27,77^. Importantly, our work extends these prior imaging observations to the full diversity of HSPC states. Moreover, by providing a more complete measure of the BM signaling environment, we show that compositionally distinct anatomical niches converge on similar HSPC-supportive signaling programs.

It is important to recognize that this distributed support model does not imply that individual cell–cell interactions are unimportant for HSPC fate. Rather, it suggests that core HSPC-supportive functions can be assembled from overlapping signals provided by multiple cellular sources in distinct anatomical environments. Indeed, this model is compatible with numerous perturbation studies demonstrating that individual niche-derived signals can influence HSPC behavior. For example, Kitl is supplied to HSCs by both endothelial and LepR+ perivascular stromal cells and deletion from either population impairs HSC maintenance, whereas combined deletion produces a substantially greater depletion of the HSC pool^78^. Megakaryocyte-derived Cxcl4 regulates HSC quiescence, and loss of Cxcl4 significantly increases HSC proliferation and reduces self-renewal capacity yet does not fully eliminate long-term HSCs from the pool^14,89^. Similarly, endothelial-derived Dll4 restrains premature myeloid differentiation, and endothelial-specific deletion induces myeloid skewing without abrogating hematopoiesis^31^, perhaps, because Dll4 is also produced by osteoblasts^90^.

Likewise, physiological and pathological remodeling of specific niche components can substantially alter hematopoietic output. During aging, stromal IL-1 signaling promotes chronic myeloid-biased hematopoiesis^91^; breast tumors remotely reprogram marrow stromal cells to favor myelopoiesis^36^; and in hematologic malignancy perturbations of specific niche components can promote expansion or transformation of specific hematopoietic populations^92,93^. Rather than conflicting with a distributed support model, these findings illustrate its central principle: individual niche interactions can exert substantial effects on HSPC behavior without being solely responsible for the broader supportive function of the BM.

Moreover, these perturbation studies and disease mechanisms illustrate a potential advantage to a distributed support architecture. Niche-specific orchestration of early hematopoiesis would place entire hematopoietic lineages under the control of specific cell-cell interactions in defined niches; thus, loss of individual cell types or ligands in a single niche could abolish entire lineages. By contrast, in a distributed support model such perturbations might abrogate specific fate decisions in the given perturbed niche. However, that lineage would not be lost as HSPCs in different niches would still receive the proper signals.

Finally, it is also important to note the limitations of our measurements and their consistency with a distributed support model. First, while we identified two transcriptionally distinct HSC populations, which differ along the transcriptional axis associated with a long-term to short-term HSC transition, estimates of true long-term HSC numbers in the murine BM suggest that they are likely rare enough that we did not image appreciable numbers^28^. Thus, it is strictly possible that specific anatomical niches are disproportionately important for long-term HSC maintenance although these cells would need to be a small fraction of the HSC 1 population we observe to be consistent with this model^21,24,25^. Second, a diverse set of combined scRNAseq, FACS, and functional studies have demonstrated that the connection between transcriptional identity and functional hematopoietic output is not strictly one-on-one for HSPCs, and individual cells that are transcriptionally classified as the same progenitor state may, nonetheless, ultimately adopt different functional fates^30,31^. Thus, it may be that HSPC functional fate is more closely tied to niche occupancy than our transcriptomic data would suggest. Though if true, such a model would require very modest if any transcriptional differences between cells that adopt different functional fates—at least among the genes we characterized. Finally, it has been recognized that HSPCs can, in part, be mobile cells^94,95^, yet our measurements are static snapshots. Thus, it is possible that fate-determining signals are preferentially provided by specific niches but that HSPC mobility is what gives rise to the lack of a clear HSPC subset preference for individual niches. Nonetheless, the largely redundant signaling environments we infer for these niches would argue against this possibility.

Together, our findings support a model in which HSPC fate decisions are not spatially orchestrated in the BM. HSPC lineage-priming is not strongly coupled to anatomical niche location, and even profound age-associated remodeling occurs broadly across pre-existing niches rather than through formation of unique aged anatomical compartments. By separating the anatomical identity of a niche from the molecular functions provided within it, this study provides a framework for understanding how distributed microenvironmental support maintains hematopoiesis during homeostasis and how coordinated disruption of that support may contribute to aging and hematologic disease. More broadly, this model reconciles seemingly conflicting observations across the niche literature, illustrates that important cell–cell interactions need not define uniquely instructive anatomical niches, and, in the context of hematologic disease, may suggest a refocus from specific niches to the BM as a whole.

## Acknowledgments

We thank members of the Moffitt and Cantor laboratories for helpful discussions. Portions of this research were conducted on the O2 High Performance Computer Cluster, supported by the Research Computing Group, at Harvard Medical School. This work was funded through National Institutes of Health (NIH) (R01HL159106), G. Harold and Leila Y. Mathers Foundation, and Harvard Stem Cell Institute grants to J.R.M. and A.B.C.; and NIH 5T32HL007574-36 which supported C.A.R-L., and by the Charles A. King Postdoctoral Research Fellowship Program from the Bank of America Co-trustees to P.C.

## Contributions

C.A.R-L., A.B.C., and J.R.M. conceived the project. C.A.R-L. performed experiments. C.A.R-L., P.C., H.Z., and J.R.M. conceptualized and validated the RIPIN methodology. C.A.R-L. analyzed data. C.A.R-L., A.B.C., and J.R.M. wrote the initial manuscript draft. C.A.R-L., P.C., H.Z., A.B.C., and J.R.M. revised the manuscript draft. A.B.C. and J.R.M. provided supervision and funding.

## Materials and Methods

### Animals

C57BL/6J mice (JAX000664) were purchased from The Jackson Laboratory at the specified age (8, 40, or 90 weeks) and sex (male or female). Mice were housed in the Harvard Center for Comparative Medicine (HCCM) facility for no more than 3 days between receipt and tissue harvest. All mouse experiments were performed in compliance with NIH guidelines and were reviewed and approved by the Harvard Institutional Animal Care and Use Committee (IACUC) under protocol IS00003215.

### RNA Extraction

All mice were euthanized with isoflurane (Patterson, 07-890-8115) overdose followed by cervical dislocation. Mouse femurs were dissected and quickly cleaned of excess muscle. Mouse BM for RNA Integrity Number (RIN) measurement and bulk RNA sequencing was extracted from the femur as a marrow plug as described previously^96^. In short, the femur was cut below the femoral head and a solution of 1× Phosphate Buffered Saline (PBS; Thermo, AM9625) and 20 mM EDTA (Thermo, AM9262) pre-chilled on ice was slowly injected into the distal femur until a marrow plug emerged. This plug was immediately placed in a RNAlater-analogous solution (13.6 mM EDTA, 18 mM Sodium Citrate [Thermo, J63888.AP], and 3.6 M Ammonium Sulfate [Sigma-Aldrich, A2939-1KG]; pH 5.2) or a cesium sulfate solution (3.6 M Cesium Sulfate [Chem-Impex, 10294-54-9; Thermo, 218702500]; pH 7.4). Samples stabilized in these solutions were stored at room temperature until downstream processing. Tissue was dissolved in TRIzol (Invitrogen, 15596026) and RNA was extracted using a Zymo Direct-zol RNA MiniPrep Plus kit (Zymo, 11-331PB) by following the manufacturer’s instructions. RIN was quantified with an Agilent bioanalyzer. All water used for solution and tissue preparation was RNase-free generated via reverse osmosis (Synergy UV, Millipore) with a Biopack polisher (Sigma, CDUFBI001).

### Bone Marrow Processing for RNA FISH

A fixation-compatible RNA stabilization solution was prepared in advance. This cesium sulfate RNA preservation buffer^56^ comprised 3.2 M cesium sulfate, 12 mM EDTA, 16 mM sodium citrate at a pH of 7.4, after dilution with paraformaldehyde (PFA). To fix samples, 32% w/v PFA (Electron Microscopy Sciences 15714) was added to a final concentration of 3.6%. Mice were euthanized as above, femurs rapidly removed, and excess muscle stripped. Small incisions were made at both ends of the femur to allow rapid penetration of the RNA stabilization and fixation solution into BM. Femurs were submerged in this solution within 2 minutes of euthanasia, as we found that this timeframe was essential to maintain intact RNA. Femurs not processed within 2 minutes of euthanasia were discarded. To ensure even solution distribution, tubes containing individual femurs within the preservation and fixation solution were gently inverted multiple times. Samples were fixed in this solution for 24 hours at room temperature and were kept on a slowly rocking platform to ensure complete coverage and even mixing of the fixation solution.

After primary fixation, femurs were transferred to a solution of 30% w/v sucrose (Thermo, 419760010) and 4% w/v PFA in 1× PBS for further fixation and cryoprotection. Samples were incubated in this solution with gentle rocking for 12-16 hours at 4 °C. The tissue was then placed in a cryomold containing a thin layer of Optimal Cutting Temperature compound (OCT; Sakura, 25608-930) at 4 °C and the mold was then filled with 4 °C OCT. The samples were incubated at 4 °C for 30 minutes to allow penetration of the OCT, and they were then flash frozen with an ethanol-dry ice slurry. This combination of 30-minute OCT incubation followed by flash freezing was found to improve both femoral morphology and the ease of sectioning. Both femurs from individual mice were prepared in a single cryoblock. Cryoblocks were stored at −80 °C until use.

Coverslips to hold tissue slices were prepared as described previously^49,55^. Briefly, circular 40-mm-diameter, No. 1.5 coverslips (Bioptechs, 40-1313-03193) were cleaned with a 1:1 mixture of 37% hydrochloric acid (Fisher, LC149503S) and methanol (Sigma, MC0480-1), silanized in a solution of 0.2% v/v allyltrichlorosilane (Sigma, 107778) with 0.1% v/v triethylamine (Sigma, T0886) in chloroform (VWR, BDH11090-4LG), washed with ethanol, allowed to dry for 1 day, and then stored under dry conditions using drierite (Fisher, AC219085000) for no longer than 6 months before use. Before sectioning, the top surface of the coverslip was covered with a solution of 0.1 mg/ml poly-d-lysine (Santa Cruz, sc-136156) supplemented with a 1:20,000 dilution of orange-fluorescent fiducial beads (Invitrogen, F8800) and incubated for 20 minutes. The solution was removed and the coverslip dried at room temperature for 1 hour. It was then covered with 4% w/v PFA in 1× PBS for 10 minutes. This solution was then removed, and the coverslip was briefly washed in 1× PBS three times and then allowed to air dry. Coated coverslips were stored dry at 4 °C until use and were used within 24 hours.

To section tissue, cryoblocks were warmed to −20 °C in a cryostat (Leica, CM1850) and then sectioned to 10 µm thickness using disposable blades (Leica, 14035838926). Coverslips were prechilled to the same temperature, and slices were gently transferred to the coverslip surface. To affix the slices to the coverslip, they were melted by touching the opposite side of the coverslip with a gloved fingertip and then immediately refrozen by placing the coverslip on the −20 °C metal freezing plate in the cryostat. To further improve tissue adherence, coverslips were then air dried until visibly dry (60 to 120 minutes) at room temperature. Slices were post-fixed to the coverslip by incubating them in 4% w/v PFA in 1× PBS at room temperature for 10 minutes and washed briefly three times with 1× PBS. We found no obvious correlation between increased drying time and measurement quality, suggesting that RNases were permanently inactivated by our fixation approach. Immediately after fixation, tissue was permeabilized with 70% ethanol at 4 °C for 12 hours. Samples were stored in these conditions up to one month before proceeding with subsequent staining.

Fixed frozen femurs prepared without the cesium sulfate RNA preservation solution were handled as described previously^55,68^. Briefly, the preparation protocol was identical to that described above; however, fixation was performed in 1× PBS with 4% w/v PFA at the specified temperature—4 °C or room temperature—for the specified time—3, 24, or 48 hours.

Fresh frozen femurs were also prepared following a previously described protocol^55,68,97^. After femur dissection and incision, femurs were placed in pre-chilled 1× PBS supplemented with 4 mM Ribonucleoside Vanadyl Complex (RVC; NEB S1402S) and incubated on ice for 30 to 60 minutes. Femurs were then transferred to cryomolds with OCT and rapidly frozen on an ethanol-dry ice slurry, as described above. Freezing was performed within 60 minutes of mouse euthanasia. Sectioning was performed as described above; however, residual OCT was sublimated by storing the coverslips in the −20 °C cryostat for 3 hours. Slices were then post-fixed at 4 °C with pre-chilled 4% w/v PFA in 1× PBS for 20 minutes.

Perfusion fixed femurs were treated identically to fixed frozen femurs following a standard whole mouse perfusion fixation^98^. Briefly, mice were given a lethal dose of isoflurane, the chest cavity was opened, an incision was made in the left ventricle, and an 18-gauge butterfly needle (Terumo, TE3SV-18BLK-CS) connected to a 60 mL syringe (Fisher, 14-955-461) containing 1× PBS was inserted through the right ventricle into the aorta. 1× PBS was perfused through the mouse by manual syringe pressure until all blood was removed. After the outflow was visibly clear, approximately 200 mL of 1× PBS with 4% w/v PFA was perfused through the mouse until the entire mouse was stiff, approximately 5 to 10 minutes. After perfusion fixation, femurs were dissected out and fixed in 1× PBS with 4% w/v PFA at room temperature for 3 hours in addition to the time of perfusion fixation.

### MERFISH library design and construction

To profile gene expression in the mouse BM, we built two custom panels comprising 186 or 1,021 genes. The smaller 186 gene panel was composed entirely of cell-type marker genes and was used to validate MERFISH performance and optimize sample preparation. The larger 1,021 gene panel was composed of both cell-type markers and sets of genes that cover pathways relevant for HSC maintenance and hematopoietic differentiation. In some cases, these genes were drawn from the KEGG database^99^ and were included even with no prior evidence of their expression. Notable KEGG pathways included mmu04350 (TGF-beta signaling), mmu04330 (notch signaling), mmu04390 (hippo signaling), mmu04115 (P53 signaling), mmu04014 (ras signaling), mmu04512 (ECM-receptor interaction), mmu04066 (hif-1 signaling), mmu04640 (hematopoietic cell lineage), mmu04060 (cytokine-cytokine receptor interaction), mmu04061 (viral protein interaction with cytokine and cytokine receptor), mmu04514 (cell adhesion molecule interaction), and mmu04071 (sphingolipid signaling). Putative gene sets were screened for genes with exceedingly high expression using bulk RNA-sequencing collected here.

MERFISH encoding probes for both libraries were designed using a previously described pipeline^48,55^. Briefly, 30-nt target regions complementary to the RNAs of interest were designed to fall within a T_M_ range of 65 to 75 °C, a GC content range of 0.4 to 0.6, and to have gene and isoform specificity scores of 0.75 to 1.

In some instances, the isoform specificity cuts were removed, allowing probes to hit multiple isoforms of the same gene. Target regions were allowed to overlap by 20 nt. A target of 72 regions was designed per RNA with shorter RNAs having fewer potential target regions. Any RNA that could support less than 30 target regions was discarded.

Individual RNAs were assigned barcodes from a set of 217 20-bit-long or 1,240 32-bit-long, Hamming-Distance-4, and Hamming-Weight-4 barcodes, for the 186 and 1,021 gene libraries respectively. The remaining 31 or 219 unused barcodes served as ‘blank’ barcodes that report on the false positive levels in individual measurements, and blank barcode levels were used to determine confidence in the expression of different gene sets in different cell types. A set of 32 previously described 20-nt readout sequences were used to define the 20-bit-long and 32-bit-long optical barcoding schemes^48,50^. Each 30-nt encoding region was flanked with 1 adenine spacer followed by 1 or 2 readout sequences for a total of 3 readout sequences in each assembled probe. Template molecules for the production of these probes were then designed by affixing a 20-nt primer sequence and the reverse complement of the T7 promoter.

Template DNA for these encoding probe sets was ordered as a complex oligo pool (Twist Biosciences). Encoding probes were then created from these templates using published methods^47,55^. In short, DNA template was first amplified by limited-cycle PCR (New England Biolabs [NEB], M0535L), purified (Zymo Spin V with the DNA Clean and Concentrator protocol; D4004 and C1012-50), and *in vitro* transcribed (NEB, E2050S). RNA was purified by home-made SPRI magnetic beads and converted back to DNA probes by reverse transcription (ThermoFisher Scientific, EP0753), and the RNA was removed with alkaline hydrolysis. The final probe sets were purified by SPRI magnetic beads and concentrated to 500 µM by ethanol precipitation. Probes were stored in water at −20 °C until use.

### MERFISH imaging system

MERFISH was performed on home-built microscopes and fluidics systems as previously described^49,50,55^. In short, custom inverted fluorescence microscopes were composed of a Nikon Ti2 microscope body, 60× CFI PlanApo Lambda oil-immersion objective affixed to a piezoelectric objective positioner (Mad City Labs, Nano-F200), motorized XY stage (Marzhauser SCAN IM 130×85 or Zaber ASR100B120B-T3A-K0066), and a Lumencor Celesta laser engine equipped with five-color excitation with center wavelengths at 405, 477, 546, 638, and 748 nm. Epifluorescence imaging was performed by passing light through a penta-band excitation filter (Semrock, FF01 391/477/549/639/741 25) and directing it onto the sample with a matching penta-band dichroic mirror (Semrock, FF421/491/567/659/776 Di01 25×36). The illumination area was an approximately 230 μm square region of the specimen. Fluorescent emission from the sample was split via a TwinCam image splitter (Cairn Research) utilizing a long-pass dichroic element (Chroma, T750LPXRXT UF2) that separated 750 nm or longer wavelengths from shorter wavelengths. Both channels were recorded by separate CMOS cameras (Hamamatsu ORCA Flash 4.0). Z-plane focus was maintained between sequentially imaged fields of view and hybridization rounds by a custom-built autofocus system composed of a 940 nm laser (Thorlabs, LP940 SF30) divided into two polarization components, reflected at distinct angles from the coverslip–sample interface, and projected onto a CMOS detector (Thorlabs, DCC1545M). All microscope operations, stage control, and fluid-handling routines were coordinated using custom software available at github.com/ZhuangLab/storm-control.

### MERFISH staining and imaging

MERFISH library and antibody staining was performed as previously described^49,50,55^. Briefly, the 70% ethanol was removed from samples, and they were washed once with 30% v/v formamide (VWR, IC11FORMD002) in 2× Saline Sodium Citrate (SSC; ThermoFisher Scientific, AM9765). Samples were then placed tissue side up on a parafilm piece to prevent sliding and covered with a 50 µL droplet of hybridization buffer (2× SSC with 30% v/v formamide, 1 mg/mL yeast tRNA [ThermoFisher Scientific, 15401029], and 10% w/v dextran sulfate [Sigma, S4030]), supplemented with 1 µM anchoring probe (/5Acryd/TTG AGT GGA TGG AGT GTA ATT+ TT+ TT+ TT+ TT+ TT+ TT+ TT+ TT+ TT+ T sequence; purchased from Integrated DNA Technologies, IDT), and 5 µg/mL of a monoclonal antibody against the Na^+^/K^+^-ATPase (Abcam, ab76020). Samples were incubated at 37 °C for 24 hours and then washed twice with 30% v/v formamide in 2× SSC at 47 °C for 30 minutes.

The samples were then stained with hybridization buffer supplemented with 1 µM anchoring probe and 5 µg/mL of a methacrylate-modified secondary antibody. Samples were incubated at 37 °C for 24 hours and then washed twice with 30% v/v formamide in 2× SSC at 47 °C for 30 minutes. The methacrylate-modified secondary antibody was included for immunofluorescent labeling of cell boundaries after tissue digestion and was produced as previously described^100,101^. Briefly, methacrylate NHS (Sigma, 730300-1G) was diluted to 100 mM in anhydrous DMSO (Thermo, D12345), and a goat anti-rabbit IgG labeled with Alexa488 (Thermo, A32731) was mixed with 100 mM MA-NHS at a 2:1 volumetric ratio and incubated in the dark for 1 hour at room temperature.

The samples were then embedded in a thin polyacrylamide film by washing the sample 2 times with gel solution (4% w/v 19:1 acrylamide/bis-acrylamide [Bio-Rad, 1610144] in 50 mM Tris-HCl [ThermoFisher Scientific, 15568-025], 300 mM NaCl [ThermoFisher Scientific, AM9759], 0.03% w/v ammonium persulfate [Sigma, 215589], and 0.15% v/v tetramethylethylenediamine [TEMED; Sigma, T7024]), and then gently inverting the coverslip onto 65 µL of gel solution on a Gel-Slick-(Lonza, 50640)-coated microscope slide (Gorilla Scientific, 6101) and allowing the gel to polymerize for 2 hours. The sample was then washed with 2× SSC and digested with 3 mL of a digestion buffer comprising 1:100 proteinase K (NEB, P8107S) in 0.25% v/v Triton-X (Sigma, T8787), and 2% v/v Sodium Dodecyl Sulfate (SDS; ThermoFisher Scientific, AM9823) in 2× SSC. This digestion was performed at 37 °C for 24 hours and then repeated one additional time with fresh digestion buffer. The sample was then washed with 2× SSC at room temperature for 30 minutes, and the wash was repeated five times to remove all excess SDS. Gel from the edges of the coverslip outside of the tissue area was trimmed away and the sample was then washed with 30% v/v formamide in 2× SSC. 50 µL of hybridization buffer with 20 µM of the 186-gene library or 50 µM of the 1,021-gene library was added to the top of the gel, and the sample was incubated at 37 °C for 72 hours. The sample was then washed with 30% v/v formamide in 2× SSC for 30 minutes at 47 °C, and the wash was repeated once.

MERFISH imaging was performed as previously described^49,50,55^. In short, coverslips were mounted in a Bioptechs FCS2 flow chamber, where buffer exchanges were performed using either a peristaltic or a syringe pump (Gilson MP1; Hamilton PSD4) to pull buffers and a series of automated valves (MVP or MVP4; Hamilton) to control the buffer pulled. Staining and imaging was performed by a sequential process comprising staining of readout probes, imaging, and cleavage of readout probe fluorophores. This process was performed with four buffers: a readout hybridization buffer comprised of 2× SSC supplemented with 10% v/v ethylene carbonate (Alfa Aesar, A15735 36), 0.25% v/v Triton X 100 (Sigma, T8787), and 15 nM each of the appropriate pair of readout probes; a readout wash buffer, which was identical to the readout hybridization buffer but did not contain readout probes; a cleavage buffer comprised of 50 mM tris(2 carboxyethyl)phosphine (TCEP; GoldBio, TCEP25) in 2× SSC; and an imaging buffer composed of 300 mM NaCl, 50 mM Tris HCl pH 8, 5 mM EDTA, 50 µM Trolox quinone, 0.5 mg/mL Trolox (Abcam, ab120747), 0.2% v/v rPCO (OYCAmericas, 46852004), and 5 mM protocatechuic acid (Sigma, 37580). A first round of readout probes and DAPI (Invitrogen, D1306) were stained to assess sample morphology and FISH quality on the microscope before full imaging. RNA FISH images were acquired with 748 and 638 nm excitation (Alexa750 and Cy5), fiducial bead images were acquired with 546 nm excitation, Na^+^/K^+^-ATPase immunofluorescence images were acquired with 477 nm excitation, and DAPI images were acquired with 405 nm excitation. Samples that did not display bright easily resolved puncta were discarded. Samples were incubated in readout hybridization buffer for 17 minutes, readout wash buffer for 6.5 minutes, and cleavage buffer for 18 minutes in each round.

### Quantification and Statistical Analysis

#### Bulk RNA sequencing

Bulk RNA isolated from whole femoral BM of 8-, 40-, and 90-week-old male mice was sequenced by Azenta Life Sciences/GENEWIZ utilizing their standard RNA sequencing protocols. In short, poly(A)-selected RNA libraries were sequenced on an Illumina NovaSeq X Plus using 2 × 150-bp paired-end sequencing, with a target output of approximately 30 million paired-end reads per sample. Raw sequencing reads were delivered as FASTQ files, aligned to the mouse GRCm39 reference genome (NCBI accession GCF_000001635.27), and processed with TopHat^102^. Transcript abundance was quantified from aligned reads using Cufflinks^103^ and reported as fragments per kilobase of transcript per million mapped reads (FPKM).

#### MERFISH image decoding

MERFISH images were processed with a previously established pipeline^49,50,55^. Briefly, images from separate rounds were re-registered using images of the fiducial beads, different readout intensities were normalized between imaging rounds, and individual pixels were assigned to barcodes based on their normalized intensity traces across the MERFISH imaging rounds using a soft decoding approach. Adjacent pixels assigned to the same barcode were aggregated to create final identified RNAs. Metadata associated with the decoding of each RNA—i.e., the number of pixels assigned to an RNA, their aggregate brightness, and the distance to the matching RNA barcode—were used to compute a quality-score associated with each RNA by comparison of these metadata properties observed for blank barcodes, as described previously^97^. RNAs were then filtered based on a quality-score threshold of 0.6.

#### Cell segmentation

Individual RNAs were grouped into single cells in a two-step process. First, we identified the boundaries of cells using CellPose v2.2.1^57^ trained on immunofluorescence images of the Na^+^/K^+^ ATPase antibody. Individual boundaries were identified in each z plane and then combined into single cells across z planes using the intersection over the union of the mask boundaries between z planes. RNAs were then assigned to individual cells based on these boundaries. We then refined this assignment using the identity and the distribution of these RNAs using Baysor v.0.7.1^58^ with the following parameters: min_molecules_per_gene was 1, min_molecules_per_cell was 50, min_molecules_per_segment was 20, confidence_nn_id was 20, n_clusters was 10, scale-std was 25%, and prior-segmentation-confidence was 0.8. Autofluorescence in the 477-nm channel associated with the immunofluorescence imaging challenged the identification of some cells with CellPose; however, we found that Baysor was able to identify RNA associated with these cells in most cases. Blank barcodes were included in Baysor assignment.

#### Single-cell analysis

Single cell analysis was performed using Scanpy v1.11.4^104^ and rapids-singlecell v0.15.0^105^ for gpu-accelerated processing. Initial embedding of all datasets was performed by concatenating raw counts from individually segmented datasets together. Cells were filtered for the number of transcripts per cell (between 5 and 5000 counts per cell) and estimated cell area (greater than 0.01 µm²). Transcript counts were normalized to the median total transcript count per cell using Scanpy’s normalize_total function, followed by a natural logarithm transformation of one plus the normalized counts using Scanpy’s log1p function. No batch correction or dataset specific normalization was performed. A UMAP representation of all cells was generated with built-in Scanpy functions by utilizing a jackstraw approach to determine the number of principal components for a cosine-distance nearest-neighbor graph. All genes included in the cell by gene matrix for principal component analysis, with the exception of blank barcodes and three excluded genes. Specifically, we excluded Cd3d, Cdh3, and Hes1 from all analyses presented here, as we found that their abundance deviated significantly from that expected from bulk RNA sequencing and showed substantial spatial distributions not associated with the BM.

Major cell classes, e.g., Hematopoietic Stem and Progenitor Cells (HSPC), Myeloid Cells, Lymphoid Cells, Megakaryocyte and Erythroid lineage Cells (MegEry), and Mesenchymal Cells, were identified via Leiden clustering with a resolution of 1.8. This analysis also identified clusters with high false positive rates, and cells within these clusters were removed from further analysis. Cell types and states within these classes were then identified through a second tier of clustering by generating a class specific AnnData object, generating a UMAP representation with the same normalization and principal component parameters as described for the all cell UMAP, and Leiden clustering with the following resolutions: HSPC: 1.0, Myeloid: 0.5, Lymphoid: 1.0, MegEry: 1.0, and Mesenchymal: 0.5. To generate the final UMAP representations for Myeloid, Lymphoid, and MegEry classes, the classes respective progenitor populations—Mono and Gran Prog, Lymph Prog, and MegEry Prog, respectively—were added to the class AnnData object and a final UMAP representation was generated.

#### Cell type annotations

Tier 1 HSPCs were identified by expression of canonical early hematopoietic markers, including Kit and Cd34. HSPC subtypes were annotated by integrating three complementary features: cluster-specific marker gene expression, pseudotemporal position along the HSPC differentiation continuum, and HSC- and lineage-associated transcriptional program scores calculated from genes represented in the MERFISH library. An HSPC specific normalized AnnData object, principal components, and UMAP representation were calculated as described above and used for subsequent HSPC cell annotation analysis.

For diffusion pseudotime analysis, a 10-nearest-neighbor graph was constructed using the first 10 principal components, followed by calculation of 30 diffusion components. An HSC-like root cell was determined with the Scanpy score genes function utilizing expression of Hlf, Mecom, Mllt3, Procr, Pdzk1ip1, Fgd5, Slamf1, Ly6a, Ifitm1, Cd27, Cd34, Cdk6, Etv6, Epha2, Il1r1, Gata2, Serpine2, Selp, Pbx1, Angpt1, and Meis1. Diffusion pseudotime was calculated in Scanpy using the first 10 diffusion components and the selected HSC-like root cell, with explicit branch detection disabled. We interpreted pseudotime as the transcriptional progression away from the most HSC-like cell along the HSPC manifold.

An HSC-enrichment score and lineage-priming scores were calculated using the Scanpy score genes function from the log-normalized expression matrix. Gene sets were curated from established transcriptional programs associated with HSC identity and hematopoietic lineage priming, informed by established scRNAseq HSPC signatures^30,60,106^. For each cell, scores represent the mean expression of the specified gene set relative to expression-matched control genes. HSC and lineage-associated transcriptional programs were independently scored using the following gene sets: HSC [Hlf, Mecom, Mllt3, Procr, Pdzk1ip1, Fgd5, Slamf1, Ly6a, Ifitm1, Cd27, Cd34, Cdk6, Etv6, Epha2, Il1r1, Gata2, Serpine2, Selp, Pbx1, Angpt1, Meis1], Megakaryocyte [Itga2b, Gp9, Vwf, Mpl], Erythrocyte [Gata1, Zfpm1, Gfi1b, Tal1, Btrc, Casp3], Monocyte [Irf8, Klf4, Cd14, Cx3cr1, Fcgr1, Lpl, Itgax, Il6ra, Itga1], Granulocyte [Gfi1, Cebpe, Cd33, Il1r2, Inhba, Itgb2l, Ncam1], and Lymphoid [Flt3, Il7r, Dntt, Bcl11a, Ccr9, Notch1, Tcf7, Gata3, Ebf1, Pax5, Il12a].

Integration of these metrics with Leiden clustering at a resolution of 1.0 resolved two HSC states, HSC 1 and HSC 2, multipotent progenitors (MPPs), and lineage-primed Megakaryocyte and Erythrocyte Progenitors (MegEry Progs), Granulocyte Progenitors (Gran Progs), Monocyte Progenitors (Mono Progs), and Lymphoid Progenitors (Lymph Progs). HSC 1 and HSC 2 expressed early stem-cell-associated genes including Kit, Meis1, Angpt1, Etv6, Pbx1, and Slamf1, whereas MPPs were characterized by Cd34, Cdk6, Fli1, and Itga2b. Lineage-primed progenitors expressed established lineage-associated transcriptional programs. MegEry Progenitors expressed Tal1, Gata1, Zfpm1, and Gfi1b; Lymphoid Progenitors expressed Dntt, Il7r, Ebf1, Pax5, and Ccr9; Monocyte Progenitors expressed Cx3cr1, Il6ra, Klf4, Itga1, and Fcgr1; and Granulocyte Progenitors expressed Gfi1, Cebpe, Itgb2l, and Hk3.

Tier 1 myeloid cells were marked by Hk3 and Irf8. To annotate Myeloid Leiden clusters, gene expression programs were manually aligned to literature markers. Immature and mature dendritic cell states were distinguished by Irf8, Siglech, Flt3, Sdc4, H2-Eb1, H2-DMb1, Itgax, and Tmem176b; macrophages were marked by C1qb, C1qc, Hmox1, Selenop, and Vcam1; myeloid derived suppressor cells (MDSCs) expressed Tlr4, Acvr2b, Il23a, Fgf22, and Ccr9; and Neutrophils were marked by Ncam1, Plaur, Tnfsf13b, Inhba, Il1rn, Cd33, Ccr1, Thbs1, Il1b, Cxcr1, Itgal, and Il1r2. We also captured a series of granulocytes, with basophils expressing Gata2, Prss34, Cd200r3, and Csf2rb2, and eosinophils expressing Prg2, Gata4, Gli2, and Il5ra.

Tier 1 Lymphoid cells were marked by Cd79a and Cd3e. To annotate Lymphoid Leiden clusters, gene expression programs were manually aligned to literature markers. Immature B committed cells ProB cells expressing Ebf1, Pax5, Sdc4, and Cplx2; PreB cells marked by Ccnb1, Hmmr, and Epha2; and more developed B-cell populations expressing Cd79a, Cd79b, Cd19, Cd22, Cxcr5, Fcer2a, Cr2, and MHC-II-associated genes including H2-Eb1, H2-Ob, and H2-DMb2. Long lived plasma cells (LLPCs) were defined by plasma cell genes Sdc1, Mzb1, Irf4, Tnfrsf17, and Tnfrsf13b. T cells expressed Ccr7, Cd3e, Cd3g, Cd5, Tcf7, and Cd8a, whereas NK cells expressed Ncr1, Klrk1, Prf1, Ccl5, and Tbx21.

Tier 1 MegEry cells were marked by Zfpm1 and Itga2b. To annotate MegEry Leiden clusters, gene expression programs were manually aligned to literature markers. Megakaryocytes were defined by megakaryocyte differentiation and platelet production genes Itga2b, Itgb3, Gp1ba, Gp5, Gp9, Vwf, Tubb1, Selp, and Thbs1. Erythrocyte populations were distinguished by expression of Steap3, Piezo1, Hmmr, and Rbl1 in Ery 1, followed by Cd36, Ackr1, Itga4, and 1300017J02Rik in Ery 2.

Tier 1 Mesenchymal cells were marked by Cxcl12, Col1a1, and Fabp4. To annotate Mesenchymal Leiden clusters, gene expression programs were aligned to literature markers. Mesenchymal Stromal Cells (MSC) cells expressed hematopoietic-supportive niche factors including Lepr, Cxcl12, Kitl, Pdgfrb, Fgf2, Igfbp5, Cxcl14, and Serping1, whereas MSC 1 and 2 were differentiated by higher expression of immune-associated genes including C4b, H2-Eb1, and H2-DMb1 on MSC 2. Vascular populations, while not universally considered mesenchymal, were grouped into the mesenchymal class and separated into sinusoidal endothelial cells (SECs) expressing Stab2, Kdr, Eng, and Tek, and arteriolar endothelial cells (AECs) expressing Cdh5, Pecam1, Cav1, Sparcl1, and Acta2. Osteolineage populations were identified by osteoblast-associated genes Alpl, Ibsp, Spp1, Mmp13, and Col1a1, and osteocyte-associated matrix genes including Dmp1, Col1a2, Fgfr1, Thbs4, Dcn, Fmod, and Postn. Adipocytes were identified by expression of Adipoq, Fabp4, Lpl, and Car3. Osteoclasts were also grouped with mesenchymal cells despite their known hematopoietic origins and were defined by Acp5, Ctsk, Tnfrsf11a, and Itgb3.

#### Embedding with single-cell RNAseq

To further validate our MERFISH measurements and cell type labels, we embedded the MERFISH measurements with existing scRNAseq of the murine BM. Mouse scRNAseq data were obtained from Young et al.^59^, and the published cell annotations were used. The scRNAseq and MERFISH datasets were restricted to genes measured in both datasets. Raw counts from the two modalities were concatenated and processed jointly using Scanpy^104^. Cells and genes were filtered using the same count-based criteria used for MERFISH analysis, after which counts were normalized and log1p-transformed using the same parameters as described for MERFISH data alone. As with the MERFISH data, Cd3d, Hes1, and Cdh3 were excluded from dimensionality reduction. Principal-component analysis was performed followed by Harmony2^107,108^ integration using both MERFISH and scRNAseq biological replicates as a single batch variable. A cosine-distance nearest-neighbor graph and UMAP representation were generated from the Harmony-corrected principal-component space.

Correspondence between MERFISH and published scRNAseq annotations was quantified in the integrated principal-component space. For each MERFISH cell, the five nearest scRNAseq cells were identified by cosine-distance using the first 35 Harmony-corrected principal components. Neighbor contributions were weighted by the inverse of their cosine-distance and normalized to sum to one for each MERFISH cell. Weights were then summed according to the published scRNAseq annotation to generate a probability distribution over reference cell types. Cell-type similarity scores were calculated by averaging these distributions across all MERFISH cells assigned to each MERFISH annotation. For visualization, similarity scores were normalized to the maximum score within each MERFISH cell type.

Human scRNAseq data were obtained from Bandyopadhyay et al.^27^. Published cell annotations and sample metadata were retained for downstream analyses. For comparison of MERFISH measurements with literature scRNAseq data, expression dot plots were generated directly from the raw scRNAseq count matrices. Mouse scRNAseq analyses were restricted to genes represented in the MERFISH panel, whereas human analyses used the corresponding human homologues of MERFISH-targeted mouse genes. For each annotated cell population, dot size represents the fraction of cells with detectable expression and dot color represents mean transcript abundance among cells of that population.

#### Neighborhood analysis

To profile BM cellular neighborhoods, we performed a neighborhood analysis analogous to that previously described^68^. In short, a table was built with rows as ‘home’ cells and columns corresponding with the first 15 nearest neighboring cells, effectively creating a cell-by-neighboring-cell matrix. This matrix was converted into an AnnData object with home cells classified as observations and neighbor cells as variables. This AnnData object was processed with built-in Scanpy functions. 35 principal components were used to generate a cosine-distance nearest-neighbor graph and UMAP representation. Leiden clustering with resolution of 0.2 was used to group cells in similar spatial environments into statistically distinct neighborhoods. Neighborhoods were manually annotated based on cell-type abundance and visual inspection of neighborhoods in space.

#### Contact frequency

To interrogate the composition of BM microenvironments, we calculated a cell contact frequency enrichment score with our cell by neighboring cell matrices. We first created a radial neighborhood table with cells as rows and the number of each neighboring cell type within 25 µm as columns. We then counted the total observed neighbors for each cell and computed the average over all cells of a given type. For a given central cell type (*i*) and a neighboring cell (*j*), we termed this quantity In parallel, we calculated the expected neighbors based on the abundance of all cell types, which for a given cell type (j) we termed *M_j_*. Thus, for each cell type (*i*) we computed the enrichment score with a cell type (*j*) from

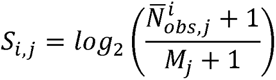

To capture the variability across different bones, 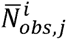 and *M_j_* were calculated for individual bones, then averaged across bones before the enrichment score was calculated.

To assess changes in cell–cell contact frequencies across age and sex, we compared the observed mean number of neighbors for each pair of cell types between biological replicates. Comparisons were performed between 8-week-old female and male mice, 40- and 8-week-old mice, and 90- and 8-week-old mice. For each cell-type pair, we added a pseudocount of 10□□ to the mean contact frequencies and calculated the log fold change between groups, retaining the direction of the change. Statistical significance was assessed using a two-sided Welch’s t-test on contact frequencies calculated independently for each bone. P values were adjusted for multiple comparisons using the Benjamini–Hochberg procedure across all cell-type pairs within each comparison.

#### HSPC niche discovery

To interrogate HSPC niches, we constructed a cell-by-neighboring-gene matrix describing the local transcriptional environment surrounding each HSPC. For each HSPC, transcripts originating from all non-home cells within a 25-µm radius were summed by gene, such that individual HSPCs were represented as observations and neighboring transcript abundances as variables. The resulting matrix was converted to an AnnData object and analyzed using Scanpy^104^. HSPCs with fewer than 100 neighboring-cell transcripts— approximately 5 percent of total HSPCs—were excluded to limit noise from sparsely sampled local environments. Counts were normalized to the median total transcript count per neighborhood using Scanpy’s normalize_total function and transformed using log1p. The 250 most highly variable genes were initially identified using the cell_ranger method, after which blank barcodes and Cd3d, Cdh3, and Hes1 were excluded from the selected features. Principal-component analysis and a cosine-distance nearest-neighbor graph were used to generate a UMAP representation of HSPC niches, and Leiden clustering at a resolution of 0.4 was used to identify recurrent niche states. Niche clusters were annotated as central marrow, sinusoidal, megakaryocytic, or endosteal based on enrichment of characteristic neighboring cell populations and inspection of their spatial localization within the BM.

To quantify the distribution of HSPC populations among these niches, niche occupancy was calculated independently for each biological replicate as the percentage of each HSPC population assigned to each niche. For a given femur and HSPC population, percentages were normalized across niches to sum to 100%; absence of that HSPC population from an individual niche was treated as 0% occupancy, whereas mice in which the HSPC population was absent entirely were excluded from the comparison. Pairwise differences in niche occupancy were evaluated using two-sided paired Wilcoxon signed-rank tests, with measurements paired by femur such that each femur was treated as a biological replicate. P values were corrected for multiple pairwise niche comparisons using the Benjamini–Hochberg false discovery rate procedure.

#### Differential gene analysis

Because MERFISH has a measurable false-positive rate arising from background signal decoded as genuine transcripts, we first estimated a comparison-specific false-positive expression threshold. This threshold was used to filter expression profiles before performing biological-replicate-level pseudobulk differential expression analyses across age, sex, and spatially defined cell populations.

To identify the false positive rate for each measurement, we leveraged the measurement of blank barcodes and the one-sided Vysochanskij-Petunin inequality as previously described^68^. In short, we calculated the probability that a given value could be contained within the distribution of our blank barcodes given their mean (μ), standard deviation (σ), and the assumption that their distribution was unimodal. We placed a 5% probability as the cut-off of our expression threshold, which equates to an expression value that is 2.81×σ greater than μ. Therefore, for each comparison, the mean and standard deviation of the raw MERFISH transcript counts associated with the 219 ‘blank’ barcodes were used to determine a comparison-specific expression threshold. Real genes with mean expression below this threshold in both the cell populations being compared were excluded from differential-expression analysis.

To perform biological-replicate-level pseudobulk differential expression analyses, a comparison-specific gene set was first determined by expression above the empirical false-positive threshold. Raw transcript counts were then summed across all cells within each biological replicate and analysis group, generating one pseudobulk expression profile per cell type or niche group for each measured femur. Pseudobulk profiles were normalized to a total of 10 counts per profile and transformed using the natural logarithm of one plus the normalized counts with the built-in Scanpy function log1p. Genes detected in fewer than two pseudobulk profiles were excluded. Differential expression was then calculated using the Wilcoxon rank-sum test implemented in Scanpy, with multiple hypothesis testing adjusted P values used to assess statistical significance.

#### Adjacent cell ligand-receptor interactions

To identify potential ligand–receptor interactions within HSPC niches, we performed a spatially restricted cell–cell communication analysis. Candidate ligand–receptor pairs were assembled from annotated interactions in the KEGG pathway database and restricted to ligands and receptors represented in the MERFISH gene panel. Analyses were performed independently for 8-, 40-, and 90-week-old mice across the four HSPC-associated niches: central marrow, sinusoidal, megakaryocytic, and endosteal. HSPC-centered niches were defined using cells and transcripts located within 25 µm of HSPCs assigned to each anatomical niche.

For cell-type-resolved interaction analysis, mean ligand and receptor expressions were calculated separately for each annotated cell population within each niche. Expression was considered above the MERFISH detection limit when the mean abundance of a transcript exceeded the mean abundance of blank barcodes plus 2.81 standard deviations within the corresponding population. Candidate interactions were retained only when both ligand and receptor exceeded this threshold in their respective populations. For each interaction, we recorded the ligand-producing and receptor-expressing cell populations, their relative abundances within the niche, and the mean log1p-transformed expression of the ligand, receptor, and any associated coreceptors. Interaction strength, reported as Combined Expression, was calculated as the product of mean ligand and receptor expression.

For calculation of aggregate niche ligand expression, the previously described cell-by-neighboring gene matrix was utilized as an aggregate of transcript abundance around HSPCs and niches were pooled by spatial location. For calculation of niche-level HSPC receptor expression, HSPCs were pooled by spatial location as opposed to cell annotation. Ligand and receptor measurements failing their respective blank-derived expression thresholds were assigned a relative expression of zero.

#### CODEX data analysis

To confirm the spatial organization of mouse and human BM, CODEX data from Li et al.^77^ and Bandyopadhyay et al.^27^ was used. Within both datasets, published cell annotations were used to define HSPC and niche cell populations. Mouse HSPCs were resolved into CD34+CD117−, CD34+CD117+, and CD34−CD117+ populations, together with common myeloid progenitors (CoMPs; lin−CD11b−Ly6C+Ly6G−) and early neutrophil progenitors (eNePs; Lin−CD117+CD71+CD49d+CD38−), providing an orthogonal view of primitive and lineage-primed hematopoietic states. Human HSPCs were resolved into HSC, SPINK2+ HSC, HSPC, GMP, CLP, CD34+CD61+, GMP/Myeloblast, and MEP/Early Erythroblast.

For each cell annotated as an HSPC, neighboring cells within a 25-µm or 50-µm radius for mouse or human, respectively, were identified using their spatial coordinates. Niche cellular composition was represented as a cell × number of neighboring cells matrix, generating an HSPC-centered representation of the local marrow environment for each cell, and converted to an AnnData object and analyzed using Scanpy^104^. Principal-component analysis and a nearest-neighbor graph were used to generate UMAP representations of HSPC niches. For the mouse CODEX dataset, Leiden clustering was performed at a resolution of 0.2. For the human CODEX dataset, counts were normalized to the median total count per neighborhood and log1p-transformed, and principal-component analysis was followed by Harmony integration across donors. A nearest-neighbor graph was constructed using the Harmony-adjusted principal components, and Leiden clustering was performed at a resolution of 0.4. Niche clusters were annotated based on enrichment of characteristic neighboring cell populations and inspection of their spatial localization within the BM.

For CODEX niche occupancy comparisons, the percentage of each HSPC population assigned to each niche was calculated independently for each biological replicate (donor or mouse). HSPC populations absent from a specific niche were assigned an occupancy of 0%, provided that the population was detected elsewhere in that replicate. For the human CODEX dataset, donors entirely lacking either HSPC population in a given pairwise comparison were excluded from that comparison. For comparisons involving the relatively rare HSC population, this criterion excluded up to five of the twelve donors. No mouse CODEX replicates were excluded for any comparison under this criterion. Pairwise differences between HSPC populations within each niche were assessed using two-sided paired Wilcoxon signed-rank tests, with biological replicates paired within each comparison. P values were adjusted using the Benjamini–Hochberg procedure independently within each niche.

