## Supplementary Information for "Early hematopoietic fate decisions are independent of bone marrow niche"

#### Supplementary Figures

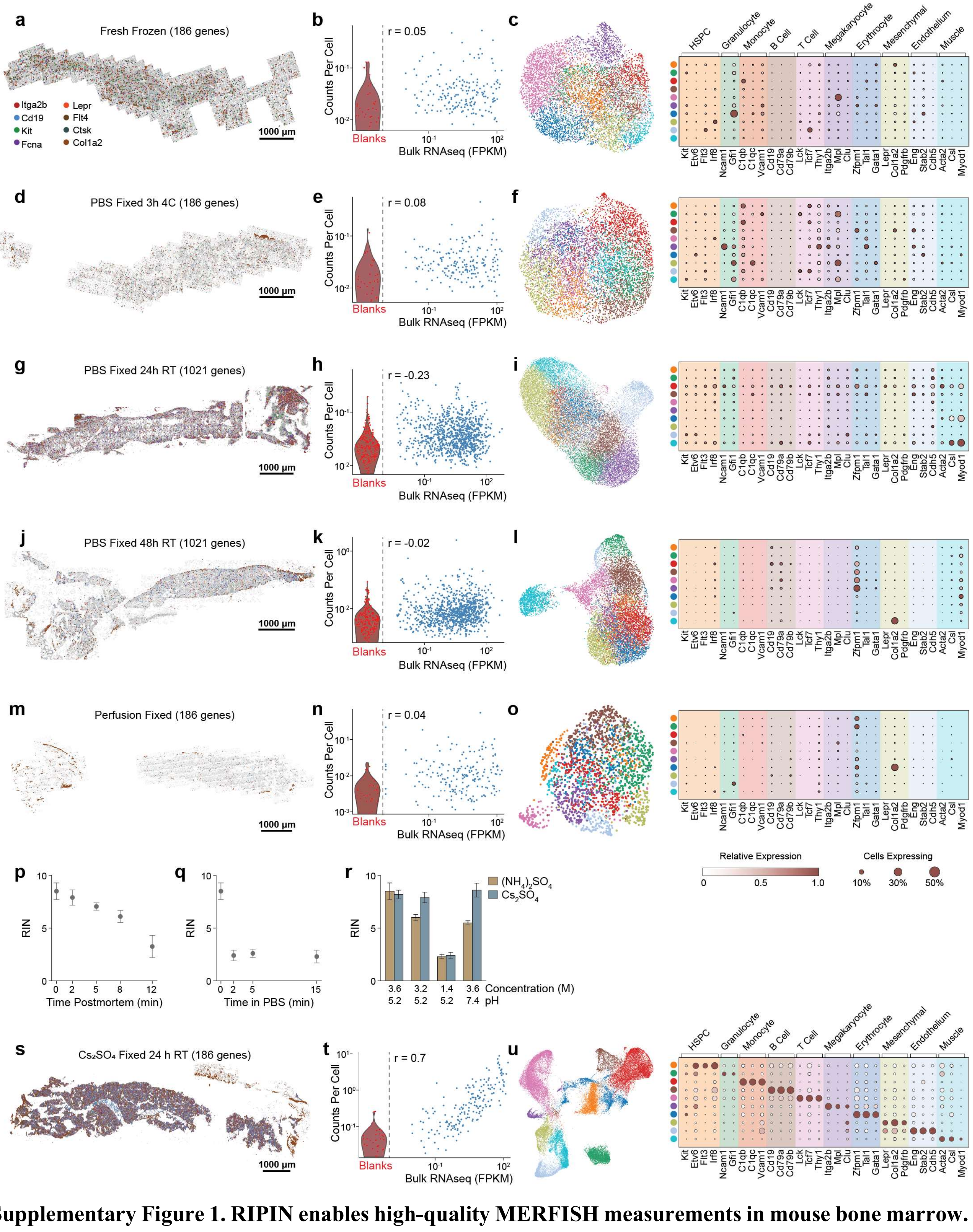


**Supplementary Figure 1. RIPIN enables high-quality MERFISH measurements in mouse bone marrow.** **a**, Spatial distribution of 8 (color) of 186 (gray) RNAs measured with MERFISH in a murine femoral bone prepared with a fresh frozen protocol. Scale bar: 1000 µm. **b**, Counts per cell for individual genes determined for MERFISH versus that determined from bulk RNA sequencing (RNAseq) of the mouse bone marrow. Histogram of the distribution of blank barcodes measured with MERFISH (red) with individual blanks shown (red markers). r: Pearson correlation coefficient between the $\log_{10}$ abundances measured with the two techniques. **c**, Left: UMAP representation of cells measured with MERFISH from the femoral bone prepared with the preparation method in (a). Color represents cell identity as determined via Leiden clustering with a resolution selected to produce 10 clusters. Right: The average expression of representative marker genes for specific bone marrow populations in each of the 10 clusters identified. Color represents the average expression while marker size represents the fraction of cells expressing at least one copy. **d-f,** As in (a)-(c) but for fixation in 4% w/v PFA in 1× PBS for 3 h at 4 °C. **g-i**, As in (a)-(c) but for fixation in 4% w/v PFA in 1× PBS for 24 h at room temperature. **j-l**, As in (a)-(c) but for fixation in 4% w/v PFA in 1× PBS for 48 h at room temperature. These data were collected with a library targeting 1,021 genes. **m-o**, As in (a)-(c) but for perfusion fixation. **p,q,** RNA integrity number (RIN) as a function of post-mortem interval (p) or time in PBS after $(NH_4)_2SO_4$ incubation (q). **r,** RIN after a 24 h incubation in candidate RNA-preservation solutions across tested concentration and pH conditions. **s-u,** As in (a)-(c) but for preparation with the RIPIN method, which involves fixation in a $Cs_2SO_4$ preservation buffer (pH 7.4) with 3.6% w/v PFA for 24 h at room temperature.

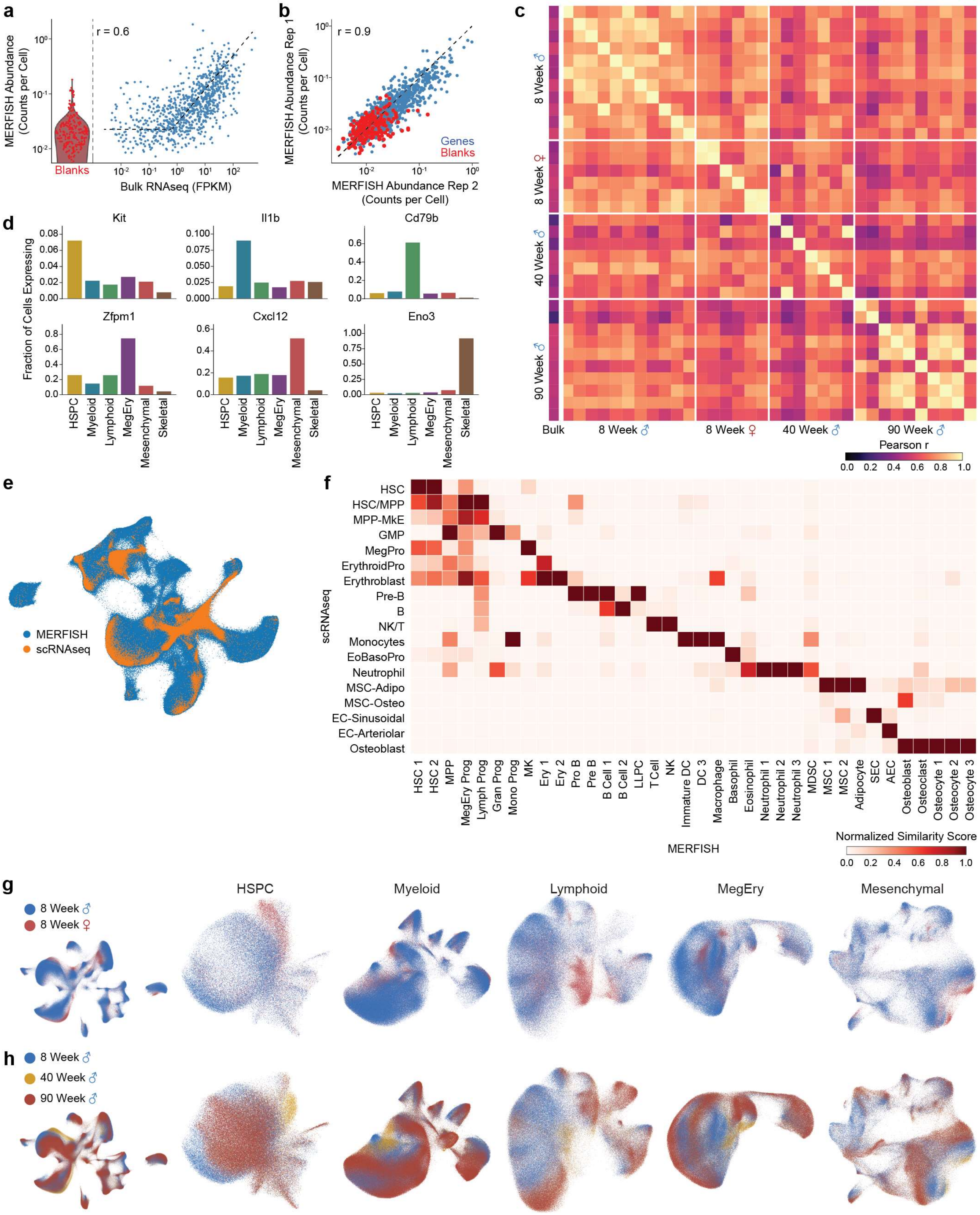


**Supplementary Figure 2. MERFISH measurements reproduce bulk and single-cell bone marrow transcriptional structure across age and sex.**

**a,** Average abundance per gene determined with MERFISH versus that determined via bulk RNAseq for an 8-week-old male bone marrow sample. The histogram represents the distribution of false-positive control barcodes (blanks) with individual controls shown in markers. r: Pairwise Pearson correlation coefficient between the $\log_{10}$ abundances. The dashed line is a guide to the eye. **b,** Average abundance per gene determined with MERFISH for one replicate of an 8-week-old male femur versus a different biological replicate from the same age and sex. Blank barcodes are marked in red. **c,** Pearson correlation coefficients between all pairs of MERFISH measurements (left) and against bulk RNAseq. The bulk RNAseq was collected for male mice at 8-, 40-, and 90-weeks of age with bulk correlations matched between mouse ages. **d,** Fraction of cells measured with MERFISH across all ages and both sexes expressing at least one copy of representative compartment markers across major cell classes. **e,** UMAP representation of the cells measured with MERFISH here co-embedded with cells from a published mouse bone marrow single-cell RNAseq (scRNAseq) reference dataset[1]. **f,** The normalized cell-type similarity scores between MERFISH-annotated cell populations and reference scRNAseq cell types from the co-embedding in (e). Scores represent the distance-weighted probability of assignment to each reference cell type based on the five nearest scRNAseq neighbors in the integrated principal-component space, averaged across MERFISH cells within each annotated population and normalized to a maximum score of 1 for each population. **g,h,** UMAP of all cells (left) or each compartment (right) colored by sex for 8-week-old mice (g) or age (h).

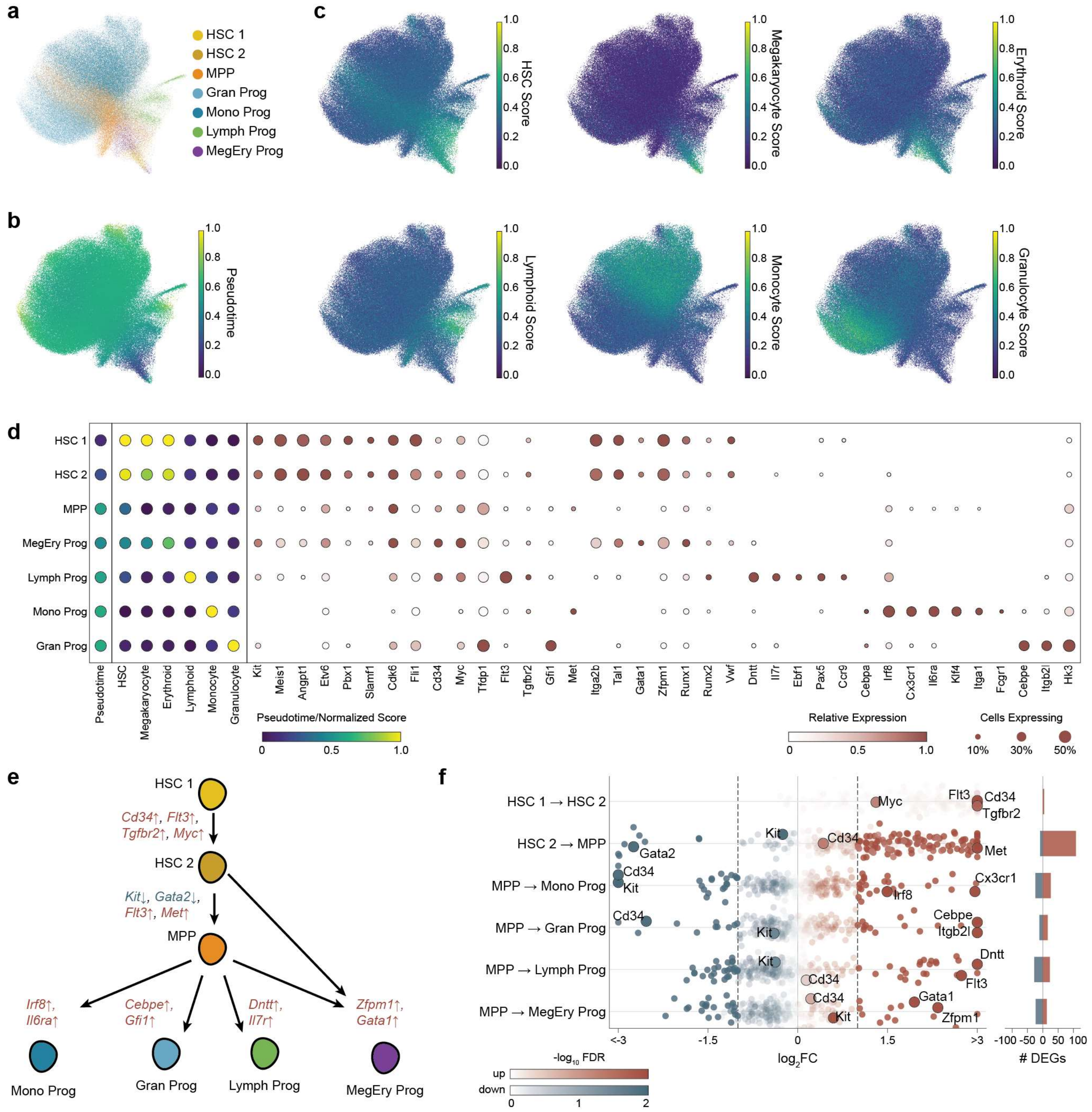


**Supplementary Figure 3. MERFISH resolves diverse and similar HSPC subsets.**

**a-c,** UMAP of HSPCs colored by cell-type annotations (a), developmental pseudotime (b), or normalized lineage scores for HSC, megakaryocyte, erythroid, lymphoid, monocyte, and granulocyte lineages (c). **d,** Average pseudotime, lineage score, or marker gene expression for different HSPC states identified with MERFISH. For gene-expression columns, size indicates the fraction of cells expressing each gene and color indicates relative expression. Dots are absent if expression was below a false discovery threshold. **e,** Schematic differentiation hierarchy from HSC 1 through HSC 2 and MPP to lineage-primed progenitors, with representative genes changing along each transition. **f,** Differential expression for all genes within each listed cell type versus that of the second listed cell type. Color indicates -$\log_{10}$ FDR-corrected P values. The side bars indicate the number of statistically significant up- (red) or down-(blue)-regulated genes as defined by an FDR-corrected P value less than 0.05 and an absolute $log_2$ fold change greater than 1. Illustrative genes are highlighted. FC: Fold change.

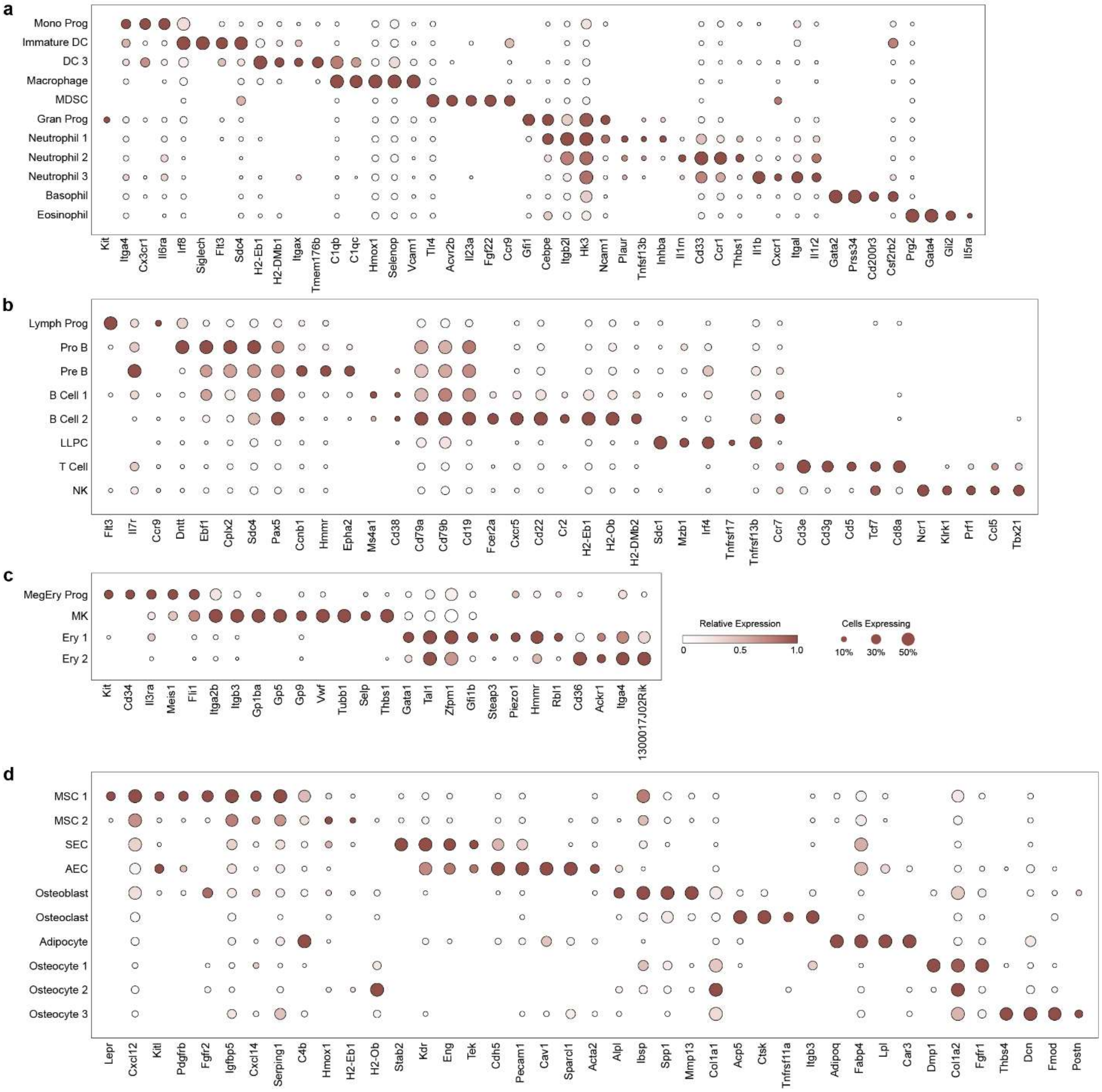

**Supplementary Figure 4. Marker expression in the identified cell populations.**

**a-d,** The average expression of marker genes across myeloid (a), lymphoid (b), megakaryocytic-erythroid (c) and mesenchymal and stromal (d) compartments. Dot size indicates the fraction of cells expressing each gene and color indicates normalized expression. Dots are absent if expression was below a false discovery threshold.

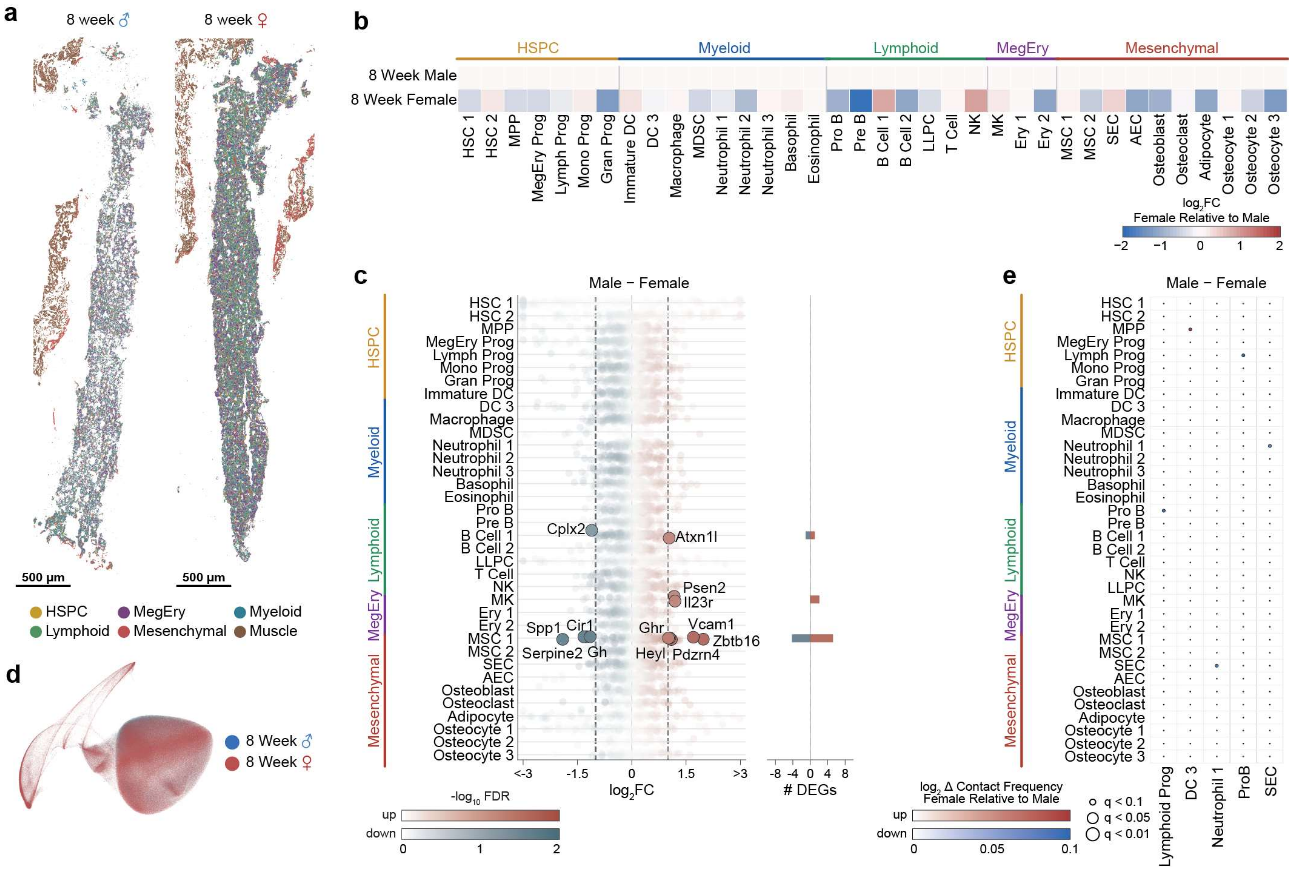

**Supplementary Figure 5. Young male and female bone marrow show limited compositional, transcriptional, and spatial differences.**

**a,** Spatial distribution of cells in representative 8-week-old male and female femoral sections colored by major cellular compartment. Scale bar: 500 µm. **b,** Cell-type abundance differences in female relative to male femurs. Relative abundance is shown as $\log_2$ fold change. FC: Fold change. **c,** Differential expression for all genes within each listed cell type between male and female mice. Color indicates $-\log_{10}$ FDR-corrected P values. The side bars indicate the number of statistically significant up- (red) or down-(blue)-regulated genes as defined by an FDR-corrected P value less than 0.05 and an absolute $\log_2$ fold change greater than 1. Illustrative genes are highlighted. **d,** UMAP representation of the cellular neighborhoods colored by sex for 8-week-old mice. **e,** Sex-associated differences in pairwise contact frequency for selected cell-type pairs (as in Fig. 2e). Dot color indicates the female-versus-male $\log_2$ fold increase (red) or decrease (blue) in contact frequency and dot size reflects the indicated FDR-corrected P values.

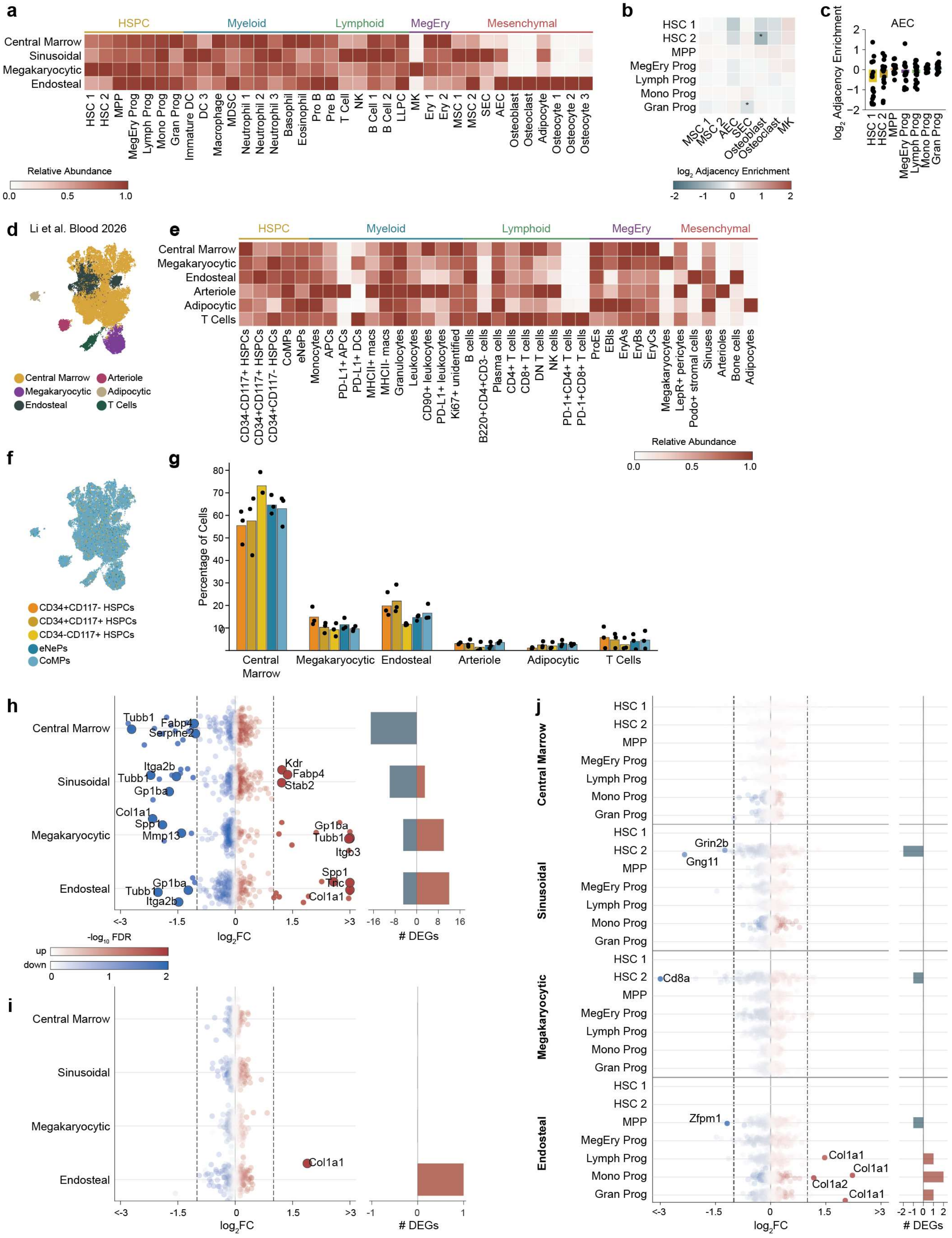

a
HSPC
Myeloid
Lymphoid
MegEry
Mesenchymal
Central Marrow
Sinusoidal
Megakaryocytic
Endosteal
HSC 1
HSC 2
MPP
MegEry Prog
Lymph Prog
Mono Prog
Gran Prog
Immature DC
DC 3
Macrophage
MDSC
Neutrophil 1
Neutrophil 2
Neutrophil 3
Basophil
Eosinophil
Pro B
Pre B
T Cell
NK
B Cell 1
B Cell 2
LLPC
MK
Ery 1
Ery 2
MSC 1
MSC 2
SEC
AEC
Osteoblast
Osteoclast
Adipocyte
Osteocyte 1
Osteocyte 2
Osteocyte 3
Relative Abundance
0.0
0.5
1.0
b
HSC 1
HSC 2
MPP
MegEry Prog
Lymph Prog
Mono Prog
Gran Prog
MSC 1
MSC 2
AEC
SEC
Osteoblast
Osteoclast
MK
log2 Adjacency Enrichment
-2
-1
0
1
2
c
AEC
log2 Adjacency Enrichment
HSC 1
HSC 2
MPP
MegEry Prog
Lymph Prog
Mono Prog
Gran Prog
d
Li et al. Blood 2026
Central Marrow
Arteriole
Megakaryocytic
Adipocytic
Endosteal
T Cells
e
HSPC
Myeloid
Lymphoid
MegEry
Mesenchymal
Central Marrow
Megakaryocytic
Endosteal
Arteriole
Adipocytic
T Cells
CD34-CD117+ HSPCs
CD34+CD117+ HSPCs
CD34+CD117- HSPCs
CoMPs
eNePs
Monocytes
APCs
PD-L1+ APCs
PD-L1+ DCs
MHCII+ macs
MHCII- macs
Granulocytes
Leukocytes
CD90+ leukocytes
PD-L1+ leukocytes
Ki67+ unidentified
B cells
B220+CD4+CD3- cells
Plasma cells
CD4+ T cells
CD8+ T cells
DN T cells
NK cells
PD-1+CD4+ T cells
PD-1+CD8+ T cells
ProEs
EBIs
EryAs
EryBs
EryCs
Megakaryocytes
LepR+ pericytes
Podo+ stromal cells
Sinuses
Arterioles
Bone cells
Adipocytes
Relative Abundance
0.0
0.5
1.0
f
CD34+CD117- HSPCs
CD34+CD117+ HSPCs
CD34-CD117+ HSPCs
eNePs
CoMPs
g
Percentage of Cells
Central Marrow
Megakaryocytic
Endosteal
Arteriole
Adipocytic
T Cells
h
Central Marrow
Sinusoidal
Megakaryocytic
Endosteal
Tubb1
Fabp4
Serpine2
Itga2b
Gp1ba
Kdr
Stab2
Col1a1
Spp1
Mmp13
Tubb1
Itgb3
Tnc
Itga2b
log2FC
### DEGs
-log10 FDR
up
down
i
Central Marrow
Sinusoidal
Megakaryocytic
Endosteal
Col1a1
log2FC
### DEGs
j
Central Marrow
Sinusoidal
Megakaryocytic
Endosteal
HSC 1
HSC 2
MPP
MegEry Prog
Lymph Prog
Mono Prog
Gran Prog
Grin2b
Gng11
Cd8a
Zfpm1
Col1a1
Col1a2
log2FC
### DEGs

**Supplementary Figure 6. HSPC anatomical niches are robustly defined in both MERFISH and multiplex proteomic imaging but do not shape HSPC transcriptional states.**

**a,** Relative abundance of cell types across central marrow, sinusoidal, megakaryocytic, and endosteal HSPC niches. **b,** Pairwise adjacency enrichment between HSPC states and niche-associated mesenchymal, endothelial, osteolineage, and megakaryocyte populations (as in Fig. 2e). *: $Log_2$ enrichments that differ from 0 with an FDR-corrected P value less than 0.05. **c,** Enrichment in the adjacency between HSPC states and AECs. Bars: mean. Markers: individual bones. No $log_2$ enrichment was statistically distinct from 0. **d,** UMAP representation of the cellular neighborhood surrounding HSPC states identified in a published multiplexed proteomic imaging dataset from healthy mouse femurs[2]. This analysis identified central marrow, megakaryocytic, endosteal, arteriolar, adipocytic, and T-cell-associated niches, which define the color. **e**, Relative abundance of cell types across the niches defined in (d). Cell type labels represent the published annotations. **f**, UMAP as in (d) but colored by the published HSPC annotations. **g**, Distribution of HSPC states across the six niches defined in (d). Bars: mean. Markers: published measurements from different femurs. **h-j,** Differential expression for all genes across all cells within a niche versus all cells in any niche (h), within all HSPC cells found in each niche relative to all HSPCs found in any niche (i), and within each HSPC state found in each niche relative to that HSPC state found in any niche (j). Color indicates $-log_{10}$ FDR-corrected P values. The side bars indicate the number of statistically significant up- (red) or down-(blue)-regulated genes as defined by an FDR-corrected P value less than 0.05 and an absolute $log_2$ fold change greater than 1. All differentially expressed genes are highlighted. FC: Fold change.

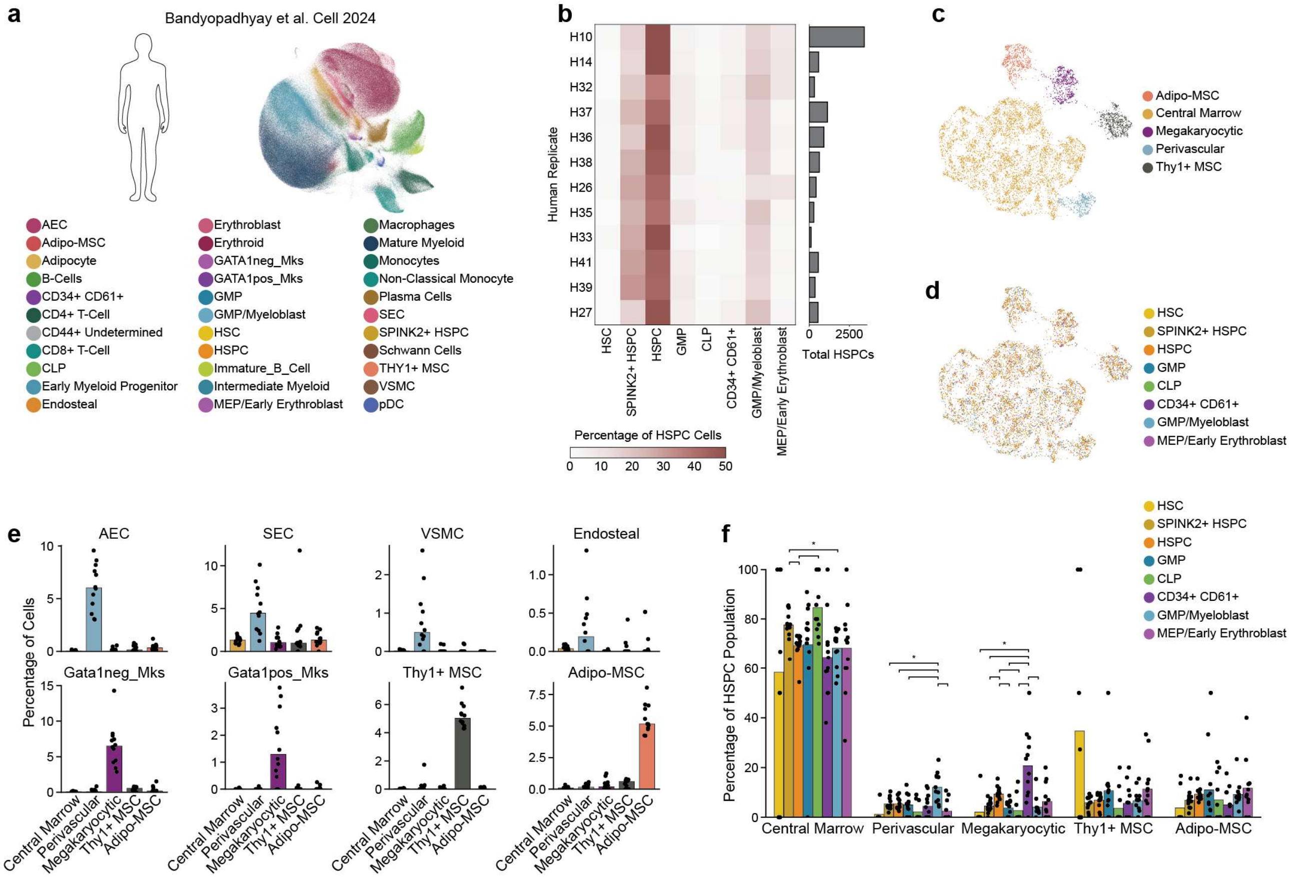


**Supplementary Figure 7. Human spatial proteomics captures anatomically defined niches and shows no niche preference for HSPC states.**

**a,** UMAP of annotated bone marrow cell states determined by multiplexed immunofluorescence and published by Bandyopadhyay et al.[3]. **b,** Composition (left) and total numbers (right) of the HSPC populations from individual human donors for the data in (a). **c,d,** UMAP of the identified cellular neighborhoods from the data in (a) colored by the niches discovered in our reanalysis (c) or by HSPC state (d). **e,** Abundance of niche-defining cell populations across the five niche classes identified in our reanalysis of the data in (a). **f,** Distribution of human HSPC states across niches. Bars: mean. Markers: donors. Brackets highlight pairwise comparisons for which the FDR-corrected P value is less than 0.05.

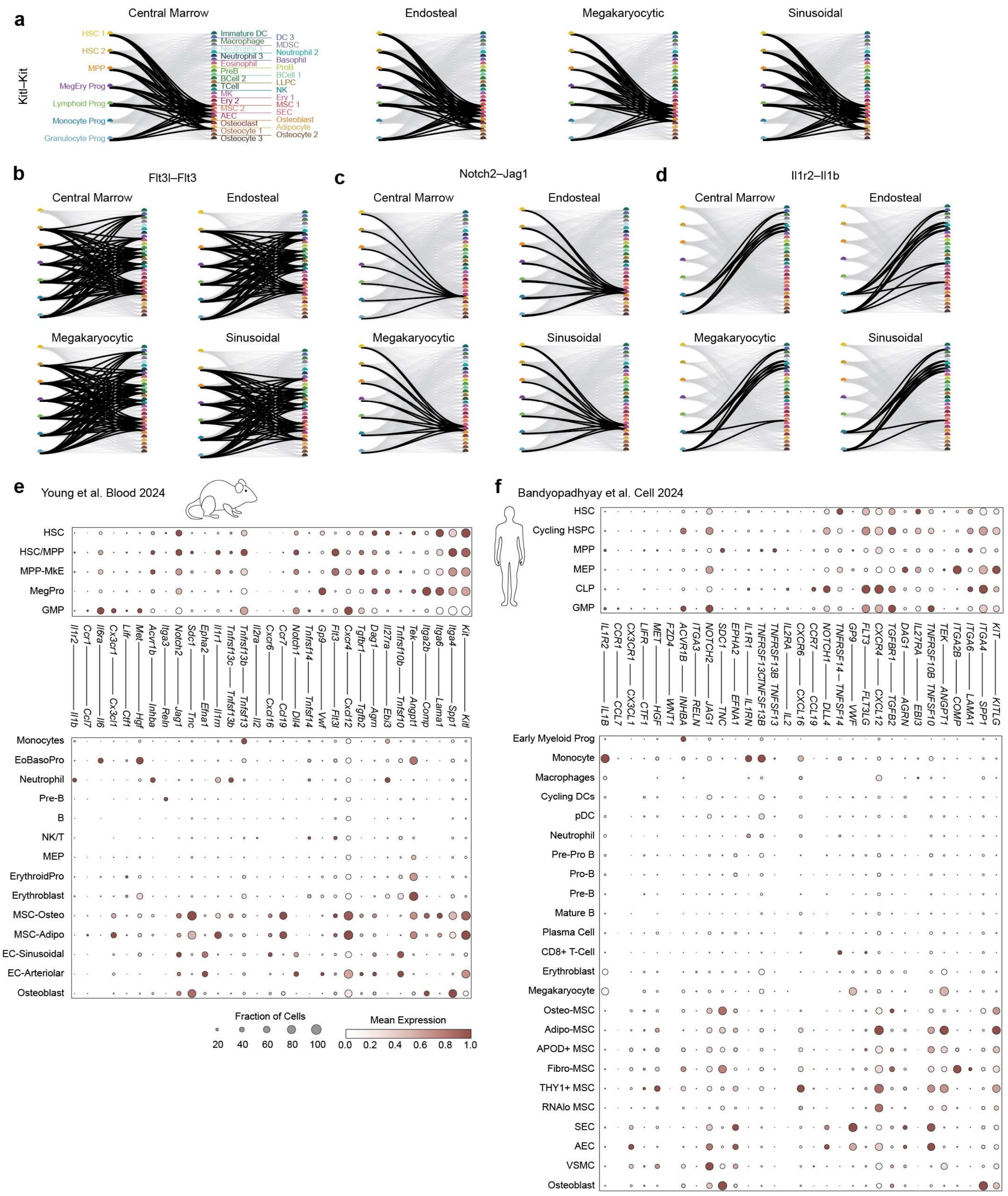


**Supplementary Figure 8. HSPC-supportive ligand-receptor interactions are distributed across cell types found in all niches and recapitulated in mouse and human scRNAseq.**

**a-d,** Sankey plots for representative ligand-receptor pathways across central marrow, endosteal, megakaryocytic and sinusoidal niches. Links connect receptor-expressing HSPC states to ligand-producing neighboring populations for Kit-Kitl (a), Flt3-Flt3l (b), Notch2-Jag1 (c) and Il1r1-Il1b (d) in each of the listed niches. All identified interactions are plotted in gray. **e,f,** Expression of the HSPC receptors and niche ligands from Fig. 3i in independent mouse bone marrow scRNAseq data[1] (e) and human bone marrow scRNAseq data[3] (f). Dot size indicates the fraction of cells expressing each gene and color indicates mean expression.

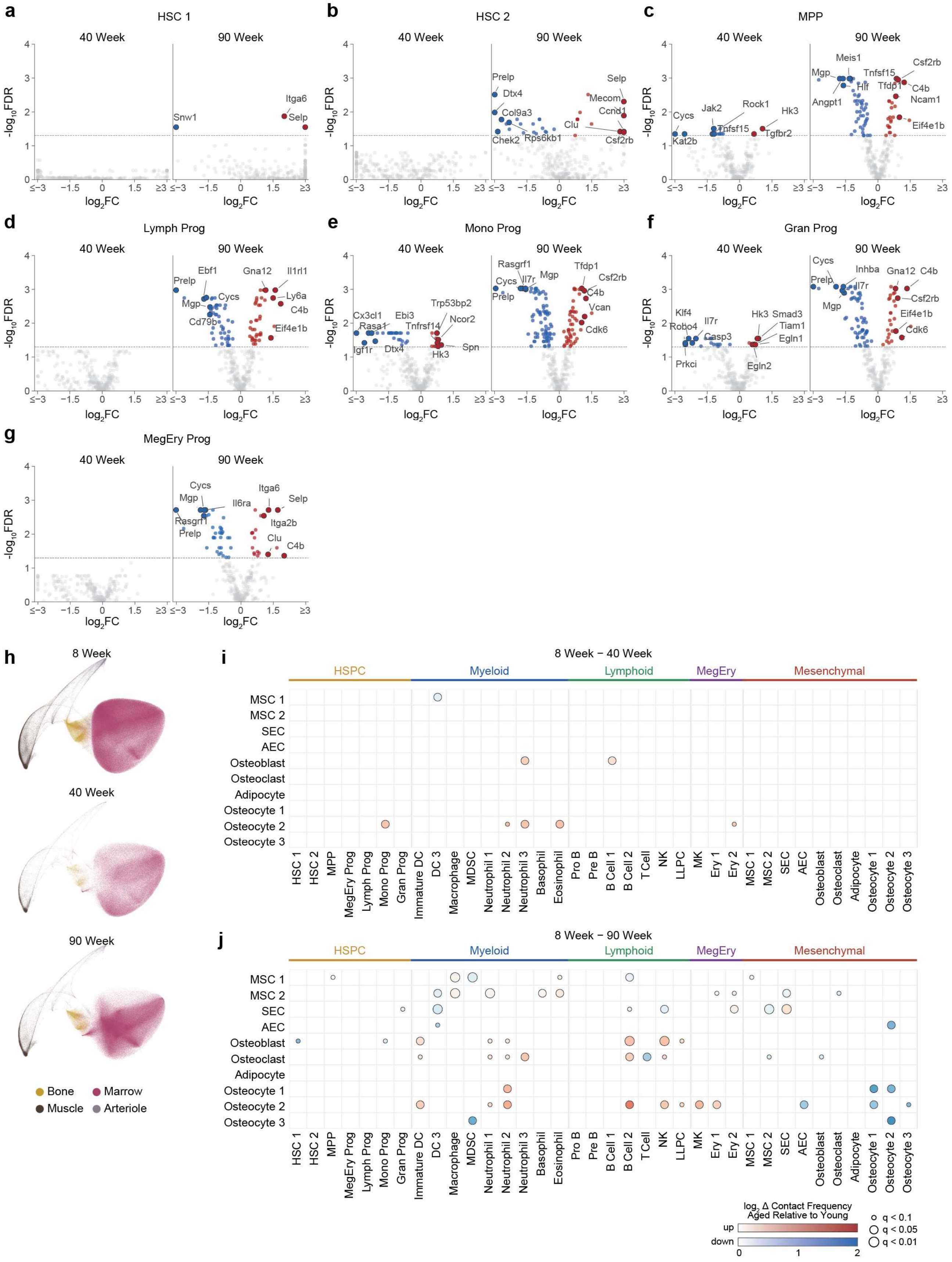

a
HSC 1
b
HSC 2
c
MPP
d
Lymph Prog
e
Mono Prog
f
Gran Prog
g
MegEry Prog
40 Week
90 Week
-log10FDR
log2FC
h
8 Week
40 Week
90 Week
Bone
Marrow
Muscle
Arteriole
i
8 Week – 40 Week
j
8 Week – 90 Week
HSPC
Myeloid
Lymphoid
MegEry
Mesenchymal
log2 Δ Contact Frequency Aged Relative to Young
up
down
q < 0.1
q < 0.05
q < 0.01

**Supplementary Figure 9. Aging progressively remodels HSPC transcriptional programs and bone marrow neighborhoods.**

**a-g,** Differential expression for all genes versus the FDR-corrected P value associated with the observed fold change for each HSPC state in the listed age versus that observed at 8 weeks. Genes are colored if they are significantly up- (red) or down-(blue)-regulated as determined by FDR-corrected P value less than 0.05. Notable genes are highlighted. FC: Fold change. **h,** UMAP of the cellular neighborhoods colored by neighborhood and containing only cells from 8-, 40-, and 90-week-old bone marrow. **i,j,** Changes in the pairwise contact frequency between niche-associated stromal populations and hematopoietic or stromal cell states at 40 weeks (i) and 90 weeks (j) relative to 8 weeks. Dot color indicates $\log_2$ fold increase (red) or decrease (blue) in contact frequency and dot size reflects the FDR-corrected significance (q).

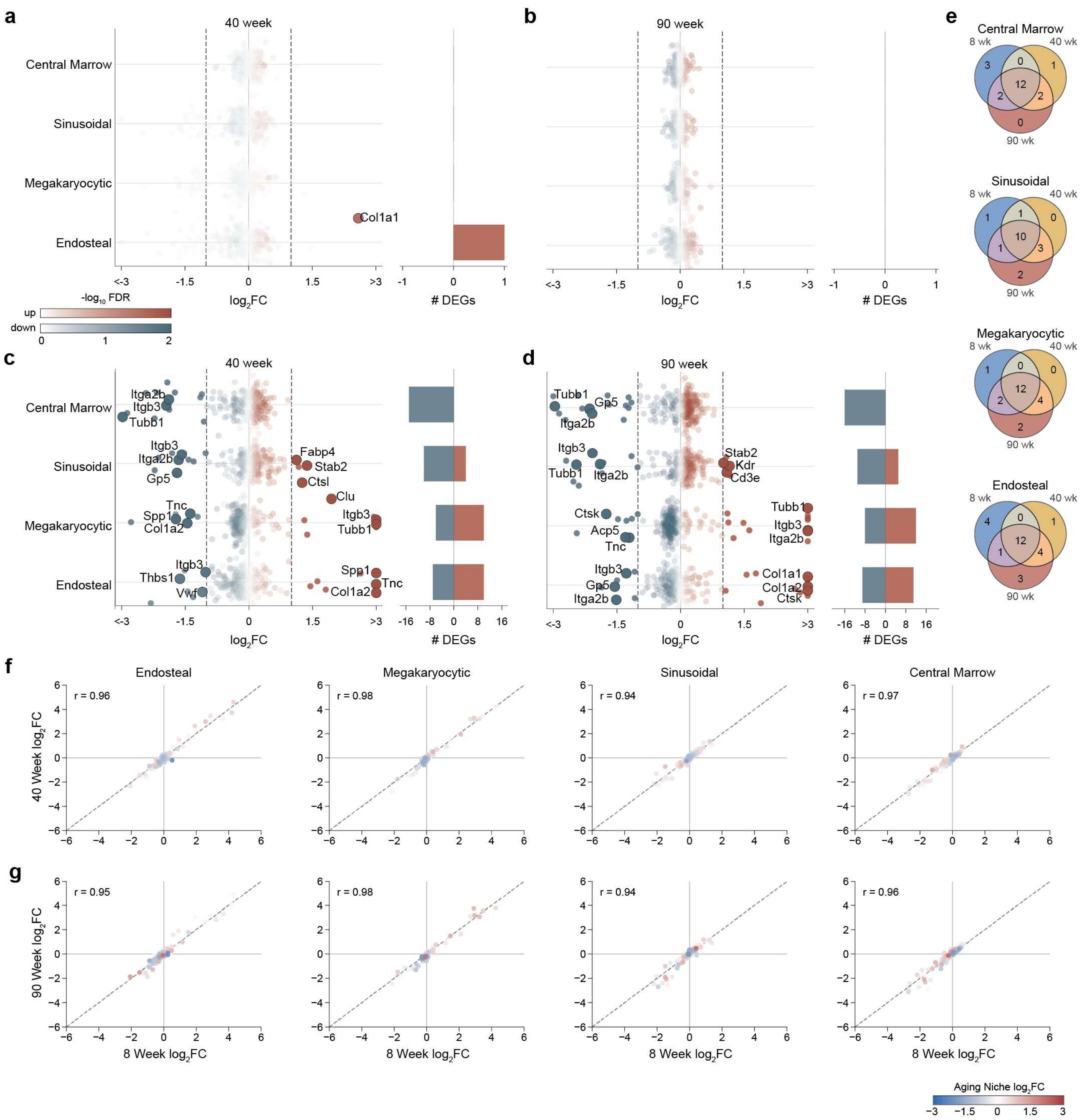

**Supplementary Figure 10. Niche-defining transcriptional profiles are conserved during aging.**

**a,b,** Differential expression for all genes across all HSPCs within a given niche versus all HSPCs seen in any niche at 40 weeks (a) or 90 weeks (b). Color indicates -$\log_{10}$ FDR-corrected P values. The side bars indicate the number of statistically significant up- (red) or down-(blue)-regulated genes as defined by an FDR-corrected P value less than 0.05 and absolute $\log_2$ fold change greater than 1. FC: Fold change. **c,d**, As in (a,b) but for all cells within a given niche versus all cells within any niche at 40 weeks (c) or 90 weeks (d). **e,** Venn diagrams of differentially expressed genes between niches, as defined for 8-week (Supplementary Fig. 6h), 40-week (c), and 90-week (d). **f,g,** The fold change in expression for each gene for all cells within a given niche versus all cells within any niche measured at 40 weeks versus measured at 8 weeks (f) or measured at 90 weeks versus measured at 8 weeks (g). The color indicates the $\log_2$ fold change for genes expressed in the listed niche at 40 weeks (f) or 90 weeks (g) versus 8 weeks. r: Pearson correlation coefficient.
